# Stabilisation of Antithetic Control via Molecular Buffering

**DOI:** 10.1101/2021.04.18.440372

**Authors:** Edward J. Hancock, Diego A. Oyarzún

## Abstract

A key goal in synthetic biology is the construction of molecular circuits that robustly adapt to perturbations. Although many natural systems display perfect adaptation, whereby stationary molecular concentrations are insensitive to perturbations, its *de novo* engineering has proven elusive. The discovery of the antithetic control motif was a significant step toward a universal mechanism for engineering perfect adaptation. Antithetic control provides perfect adaptation in a wide range of systems, but it can lead to oscillatory dynamics due to loss of stability, and moreover, it can lose perfect adaptation in fast growing cultures. Here, we introduce an extended antithetic control motif that resolves these limitations. We show that molecular buffering, a widely conserved mechanism for homeostatic control in nature, stabilises oscillations and allows for near-perfect adaptation during rapid growth. We study multiple buffering topologies and compare their performance in terms of their stability and adaptation properties. We illustrate the benefits of our proposed strategy in exemplar models for biofuel production and growth rate control in bacterial cultures. Our results provide an improved circuit for robust control of biomolecular systems.

## I. INTRODUCTION

Synthetic biology promises to revolutionise many sectors such as healthcare, chemical manufacture and materials engineering^7^. A number of such applications require precise control of biomolecular processes in face of environmental perturbations and process variability^22^. An important requirement in such control systems is *perfect adaptation*, a property whereby chemical concentrations remain insensitive to perturbations^20,23^. The molecular mechanisms that can produce perfect adaptation has been extensively studied in natural systems^3,14,20,23^. In these systems, perfect adaptation can be produced by a range of feedforward and feedback mechanisms^3,20^. Such natural systems have been shaped by evolutionary processes, but it remains unclear if they are sufficiently robust and tuneable for *de novo* engineering of perfect adaptation in synthetic circuits.

One approach to engineer perfect adaptation relies on the use of feedback control. As illustrated in Figure 1A, this strategy requires circuits that sense the output and act upon the inputs of a biomolecular process. The groundbreaking work by Briat and colleagues^1,4^ identified *antithetic feedback* as a promising candidate for engineering perfect adaptation in living systems. Antithetic control involves a feedback mechanism with two molecular components that sequester and annihilate each other (see Figure 1B). It enables a system output to robustly follow an input signal and remain insensitive to various types of perturbations, akin to what integral feedback achieves in classic control engineering strategies^2^.

**FIG. 1.**
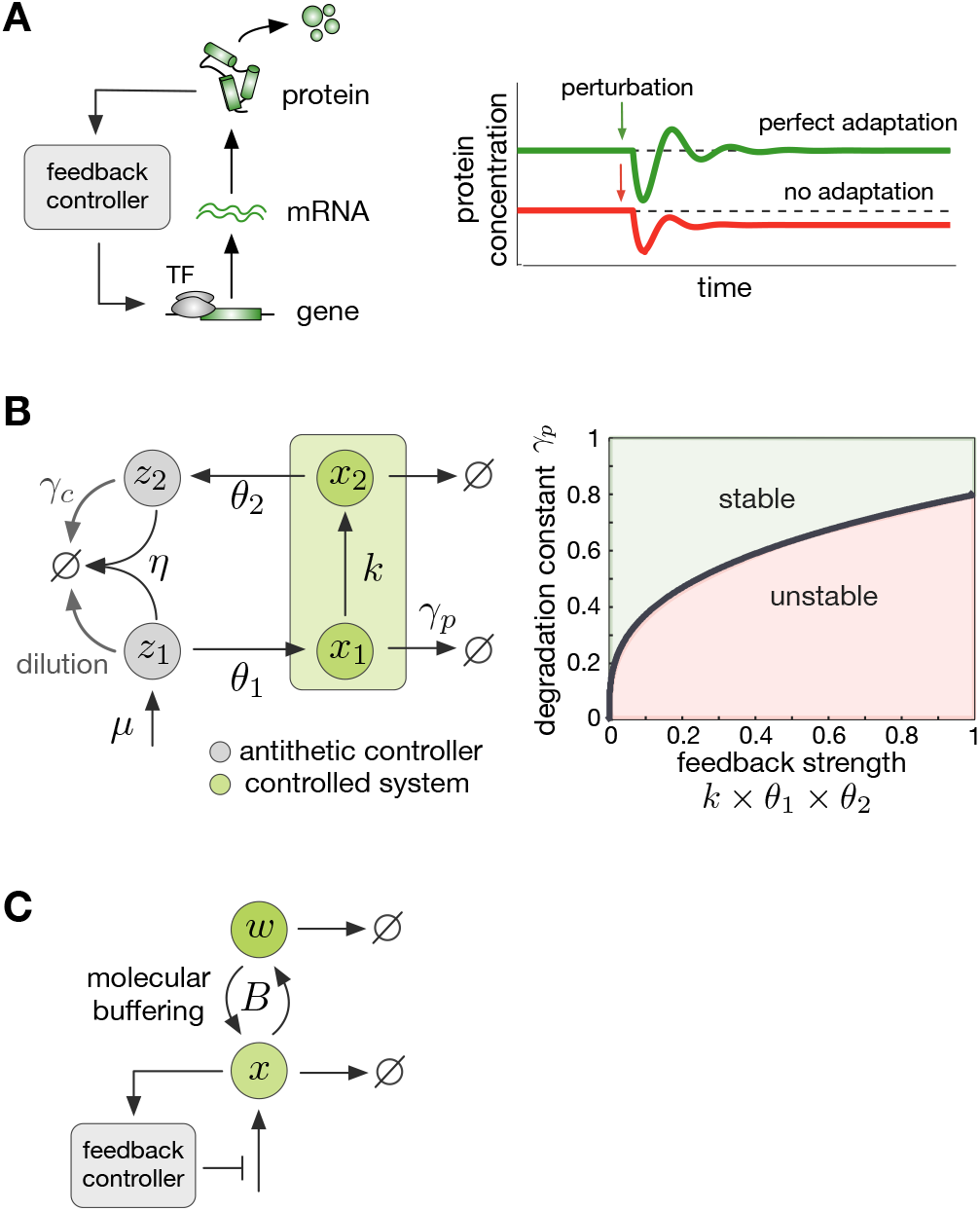
Perfect adaptation and feedback control. **(A)** Schematic of a feedback system designed to achieve perfect adaptation in protein expression. Based on readouts of protein concentration, the controller modifies the activity of a transcription factor (TF). If the controller achieves perfect adaptation, steady state protein concentrations are robust to perturbations. **(B)** Left: the antithetic feedback controller, first proposed Briat *et al* in^4^, can achieve perfect adaptation. In the presence of dilution (*γ_c_* ≠ 0), the antithetic controller does not achieve perfect adaptation. Right: conditions for stability in the case of a two-species system. The stability boundary is the condition in (2); the example was computed with fixed parameters *k* = *θ*_2_ = *γ_p_* = 1. **(C)** In the example, molecular buffering provides a general mechanism to stabilize feedback control systems^17^. The buffer reversibly sequesters molecules of species *x* into an inactive form *w*.

The original antithetic control motif, however, has two weaknesses that can limit its applicability: it is often not effective when cells are growing rapidly, and the feedback mechanism can cause unwanted oscillations under a range of conditions^27^. Specifically, dilution effects caused by cell growth cause “leaky integration” - so called because integration is a form of memory and dilution causes that memory to leak over time^29^. This prevents perfect adaptation from occurring during rapid growth. Although in some motif configurations, the loss of perfect adaptation can be partly mitigated with a stronger feedback^29^, in general the use of strong feedback results in the loss of stability and undesirable oscillations^27^. Such oscillations can be stabilised in specific motifs^15^, and in more general cases the combination of antithetic control with classic Proportional-Integral-Derivative (PID) control has been shown to improve temporal regulation^5,9^. Yet to date, there is no general strategy to avoid oscillations and prevent the loss of adaptation during rapid growth.

Here we propose an extended antithetic control system that resolves the above limitations. We show that the addition of molecular buffers improves stability and suppresses undesirable oscillations, and moreover it can allow for near-perfect adaptation in fast growth regimes. Molecular buffering is a widespread regulatory mechanism in nature (e.g. ATP, calcium & pH buffers^17,21,32^) that has received modest attention in the literature as compared to other regulatory mechanisms. Recent work found that the combination of buffering and feedback is often critical for robust regulation^16,17^. Buffering has the ability to attenuate fast disturbances and stabilise feedback control^17^, and can also be essential for the control of multiple coupled outputs^18^. Here, we first show that a number of buffering topologies can stabilise the original antithetic control system and preserve perfect adaptation. We then show that buffering can allow increased feedback strength or ‘gain’ without producing oscillations, which in turn reduces the steady state error even in fast growth conditions. To illustrate the utility of this new antithetic control strategy, we examine two case studies that involve the control of biofuel production and growth rate in microbes.

## II. BACKGROUND

### A. Perfect adaptation and antithetic control

Antithetic control employs a feedback mechanism with two molecular components that sequester and annihilate each other (see Figure 1B). In its most basic formulation, an antithetic system contains a two-species molecular process to be controlled, and a two-species antithetic controller. The two species of the controlled process (*x*_1_ and *x*_2_) can represent a variety of molecular systems, including e.g. mRNA and protein as in Figure 1A. The goal of the antithetic control system is to desensitise the steady state concentration of *x*_2_ with respect to external perturbations. Such perturbations include, for example, insults of molecular species coming from upstream or downstream processes, changes in cellular growth conditions, or alterations to binding affinities between species.

In the absence of stochastic effects, the feedback system can be modelled by the ODEs:

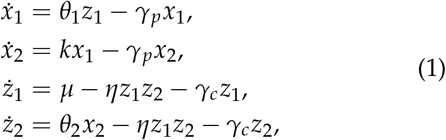

where *z*_1_ and *z*_2_ are the concentrations of species in the antithetic controller, and *θ*_1_, *k*, and *θ*_2_ are positive parameters representing first-order kinetic rate constants. The parameter *μ* describes a zero-order influx of controller species *z*_1_, while *η* is a second-order kinetic rate constant. We further assume that molecular species are diluted by cellular growth, degraded by other molecular components, or consumed by downstream cellular processes, all of which we model as a first-order clearance with rate constant *γ_p_*. The controller species *z*_1_ and *z*_2_, on the other hand, are assumed to be diluted by cellular growth with a rate constant *γ_c_*.

In the absence of dilution effects (*γ_c_* = 0), from the model in (1) we can write:

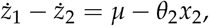

which after integration becomes

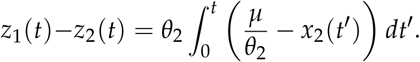

The above equation means that, if the system has a stable equilibrium, the steady state concentration of *x*_2_ is *x*_2_ = *μ* /*θ*_2_, and hence independent of all model parameters except *μ* and *θ*_2_. Therefore the antithetic control system displays perfect adaptation because the steady state of *x*_2_ is robust to perturbations in parameters *k, θ*_1_, *θ*_2_ and *η*.

A caveat of antithetic feedback is that it can have a destabilising effect. When the controller species are not diluted (*γ_c_* = 0), it can be shown that a parametric condition for stability is^27^:

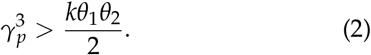

As shown by the stability diagram in Figure 1B, strong antithetic feedback can cause the system to lose perfect adaptation and display oscillatory dynamics. Moreover, in the presence of dilution of the controller species (*γ_c_* > 0), the antithetic controller is unable to produce perfect adaptation^27^. The adaptation error can be reduced with stronger feedback, for example by increasing the rate constants *θ*_1_, *k*, or *θ*_2_. Yet as mentioned above, stronger feedback can cause unwanted oscillations^27^. These caveats are particularly relevant in bioproduction applications that require fast culture growth^10,35^.

### B. Molecular buffering

Buffering is the use of molecular reservoirs to maintain the concentration of chemical species^17^. It is a widespread regulatory mechanism found across all domains of life, with common examples including pH, ATP and calcium buffering^21,32^. Molecular buffering can have a number of regulatory roles^17,18^, including acting as a stabilising mechanism for other molecular feedback systems^17^.

To provide a background on buffering models, we consider the simple case of a chemical species (*x*) that is subject to feedback regulation, as shown in Figure 1C. A general model for such process is:

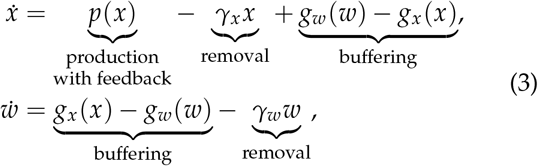

where *w* is a molecular buffer for the regulated species *x*, and *p*(*x*) is a feedback-regulated production rate of *x*. The parameters *γ_x_* and *γ_w_* are first-order clearance rate constants of *x* and *w*, respectively. The terms *g_w_* and *g_x_* describe the reversible binding of species *x* and the buffer *w*. The steady state 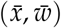 occurs when production matches degradation (i.e. 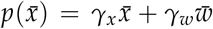) and conversion from *x* to *w* matches the reverse conversion plus removal (i.e. 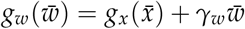).

It can be shown that after linearisation and assuming that the buffering reactions rapidly reach quasiequilibrium, the model (3) can be simplified to (see SI1):

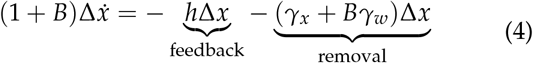

where 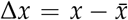 is the deviation of *x* from the steady state 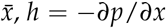, *h* = −*∂p*/*∂x* is the linearised feedback gain and *B* is the buffer equilibrium ratio:

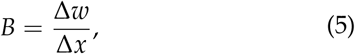

where 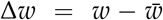 is the deviation of *w* from the steady state 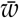. The parameter *B* is buffer-specific and quantifies the change in the concentration of a regulated species (*x*) to the change in the concentration of a buffering species (*w*) when the buffering reactions are at quasi-equilibrium^17,18^.

From (4) we observe that buffering slows down the rate of change of the output *x* by a factor of (1 + *B*). It can be shown^17^ that this slowed rate generally helps to attenuate fast disturbances and stabilise unwanted oscillations (see SI1). More generally, the stabilisation effect of buffering results from two properties. First, the buffering reactions counteract changes to a target molecular species by acting directly on the species and not via indirect or complex feedback loops^16,17^. Second, buffering can be shown to be mathematically equivalent to popular feedback strategies known to have useful stabilisation properties^2^. In particular, rapid buffering without degradation (*γ_w_* = 0) is equivalent to negative derivative feedback^17^:

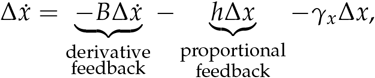

where the buffer equilibrium ratio *B* corresponds to the derivative feedback gain commonly employed in control engineering. Likewise, the general case of non-rapid buffering with degradation is mathematically equivalent to the so-called “lead” controller employed in control engineering^16^. In the next section, we study the ability of buffering to stabilise oscillations in antithetic control systems.

## III. BUFFERING CAN STABILISE ANTITHETIC INTEGRAL FEEDBACK

In this section we study a modified version of the antithetic feedback controller that includes buffering of its molecular components. We consider a number of architectures (Figure 2A) and identify those that suppress oscillations caused by the instability illustrated in Figure 1B. We show that rapid buffering without degradation does not improve stability, while non-rapid buffering and rapid buffering with degradation are highly effective stabilisers.

**FIG. 2.**
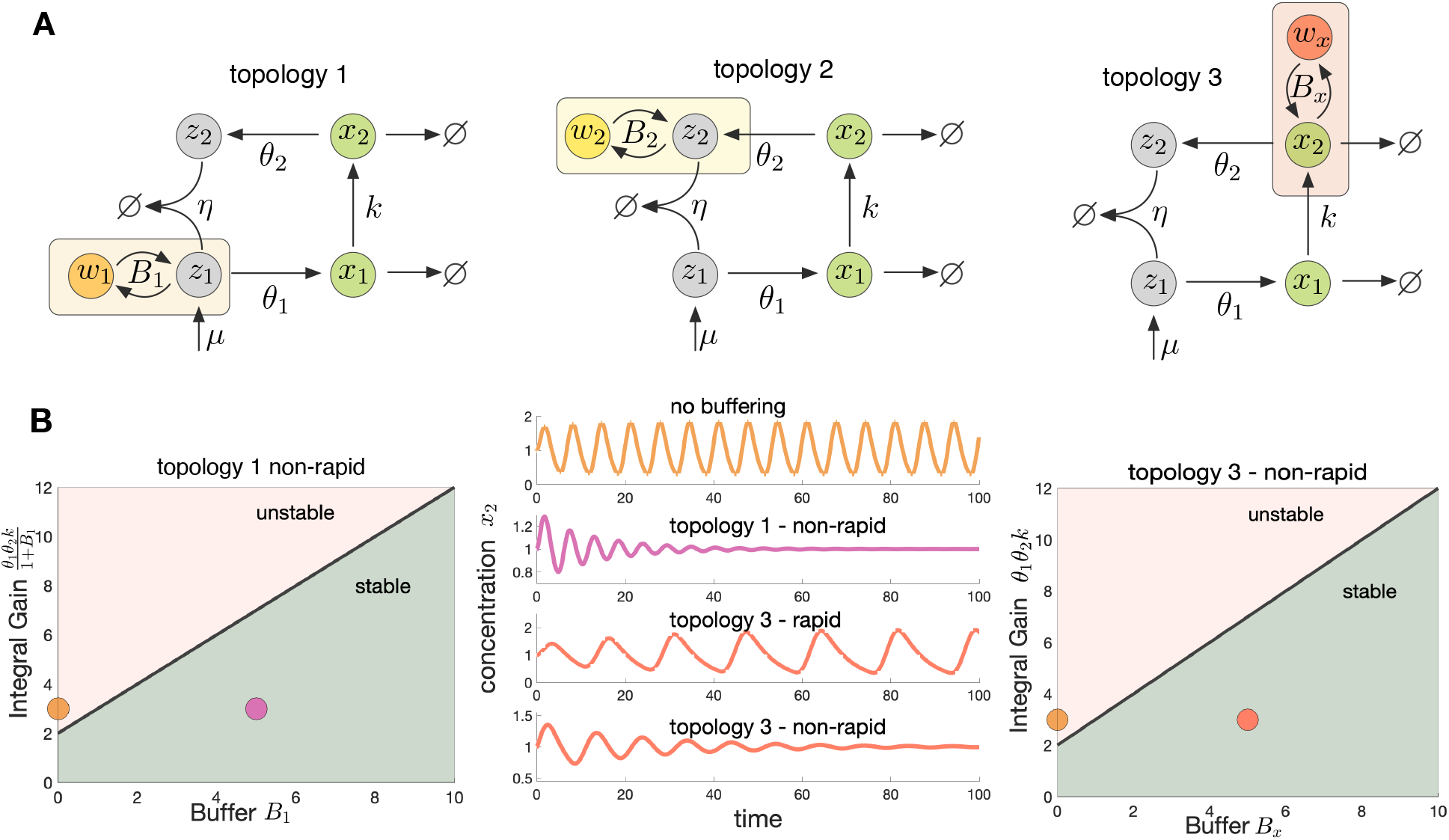
Buffering stabilizes antithetic feedback. **(A)** Schematics of three buffered antithetic systems, without dilution (*γ_c_* = 0); each topology includes buffering a specific molecular species. **(B)** Stability diagram for buffered topologies 1 and 3. The stability boundary on the left corresponds to the condition in (10) and the boundary on the right to (11). The time courses show simulations of the output species (*x*_2_) for different topologies. Parameters are *μ* = 1, *θ*_1_ = 3, *θ*_2_ = 1, *γ_p_* = 1, *η* = 100 and *k* = 1. With topology 1, parameters are *B*_1_ = 5, *θ*_1_ = 18 and *b_wz_* = 1 (non-rapid). With topology 3, parameters are *B_x_* = 5, *θ*_1_ = 3, *b_wx_* = 1 (non-rapid) and *b_wx_* = 20 (rapid).

### A. Rapid Buffering

To study the impact of buffering on the antithetic controller, we consider mathematical models for the topologies in Figure 2A, in which species *z*_1_, *z*_2_, and *x*_2_ are buffered by molecules *w*_1_, *w*_2_, and *w_x_*, respectively. For simplicity, we use linear buffering reaction rates in order to focus on the nonlinearity due to the mutual annihilation of controller species *z*_1_ and *z*_2_^9^. As in Eq. (4), we assume that the buffers rapidly reach quasi-equilibrium to obtain the following extended model (see SI2.1):

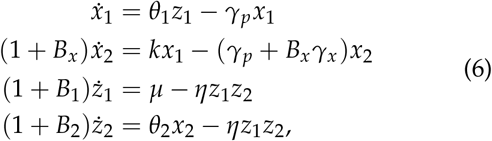

where *B_x_*, *B*_1_ and *B*_2_ are the equilibrium ratios for each buffer, and *γ_x_* represents the degradation rate of buffer *w_x_*. The extended model in (6) reduces to the original antithetic system in (1) if *B_x_* = 0, *B*_1_ = 0 and *B*_2_ = 0.

In the extended antithetic controller with rapid buffering, the parameters (*B_x_*, *B*_1_, *B*_2_, *γ_x_*) are additional tuning knobs that can be used to shape the closed-loop dynamics. In Figure 2A we show the three considered buffering architectures.

We first show that buffered antithetic feedback preserves perfect adaptation. From (6) we write

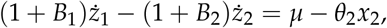

which after integrating becomes:

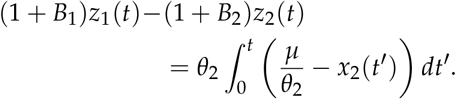

The above integral ensures that if the system is stable then the steady state of *x*_2_ is *x*_2_ = *μ* /*θ*_2_, hence independent of all parameters except *μ* and *θ*_2_. The steady state of *x*_2_ is thus robust to perturbations in the original parameters *k*, *θ*_1_, *θ*_2_, and *η*, as well as the additional parameters introduced by the buffering mechanism *B_x_*, *B*_1_, *B*_2_, and *γ_x_*. This means that the buffered antithetic feedback displays perfect adaptation as in the original formulation in Eq. (1).

Assuming strong integral binding (large *η*) then *z*_2_ is small and so with rapid buffering we have the antithetic input in to the controlled system (see Figure 1B and SI4.1)

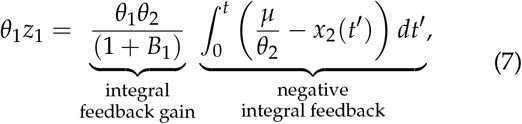

where *θ*_1_ *z*_1_ represents the control of *x*_1_ production in (6) and we can observe that the integral gain is dependent upon *B*_1_. This changes the gain because rapid buffers tend to slow down the dynamics of *z*_1_ in (6) and for integral feedback the gain is inversely proportional to the time scale of *z*_1_.

As shown by the stability condition in (2), the original antithetic control system becomes oscillatory when the feedback gain is too strong or the degradation of *x*_1_ and *x*_2_ is too slow^27^. To analyse the stabilising role of buffering, we first consider the system in the absence of degradation of the *w_x_* buffer (i. e. *γ_x_* = 0). Assuming rapid buffering and strong integral binding (large *η*), we find that the system is stable when (see SI2.1):

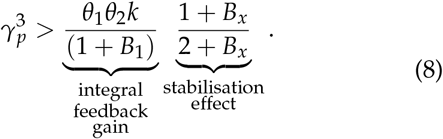

From the condition (8) we observe that increasing *B*_1_ reduces the lower bound for *γ_p_* and improves stability. However, from (7) we also observe that such effect is equivalent to a reduction in the feedback gain by a factor (1 + *B*_1_). Therefore a similar effect can be obtained simply by changing the gain *θ*_1_*θ*_2_ without the additional complexity of buffering *z*_1_. Moreover, the condition in (8) also shows that rapid buffering of *z*_2_ has no impact on stability, because the stability boundary is independent of *B*_2_, whereas rapid buffering of *x*_2_ can destabilise the system and produce oscillations, because the stability boundary becomes more stringent for large *B_x_*.

### B. Non-Rapid Buffering in Topologies 1 and 3

We next sought to determine the stabilisation properties of non-rapid buffering, i. e. implemented with buffers that do not satisfy the rapid equilibrium assumption. We show that non-rapid buffering can be a highly effective stabilisation strategy for Topologies 1 and 3, and this stabilisation effect is greater than the effect attributed to the change in feedback gain alone.

To model the effect of non-rapid buffering, we replace the model for *ẋ*_2_ and *ż*_1_ in (6) by

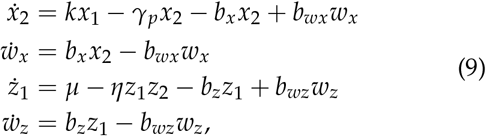

and keep the models for *ẋ*_1_ and *ż*_2_ identical as in (6). Given that in the previous section we found that topology 2 does not alter stability, from here onward we set *B*_2_ = 0. For topologies 1 and 3, the buffer equilibrium ratios (*B*_1_, *B_x_*) determine the concentration of the buffering species at equilibrium and the reverse reaction rate constants (*b_wz_, b_wx_*) determine the speed at which the buffers reach equilibrium. These four parameters can be regarded as tuning knobs for modifying the closed-loop dynamics, akin to the role of *B*_1_ and *B_x_* in the rapid buffering case of the previous section.

For Topology 1 (i.e. *b_x_* = *b_wx_* = 0) and assuming strong antithetic binding (large *η*), the stability condition is:

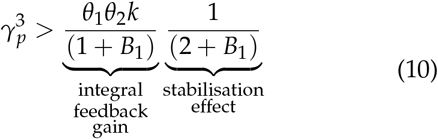

whereas for Topology 3 (i.e. *b_z_* = *b_wz_* = 0) the stability condition becomes:

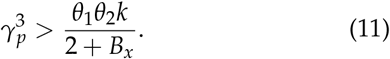

To derive both conditions (10)–(11) we have assumed for simplicity that *b_wz_* = *γ_p_* and *b_wx_* = *γ_p_*, respectively. The general stability condition for other values of *b_wz_* and *b_wx_* can be found in SI2.2-2.3.

As shown by the stability diagrams in Figure 2, the conditions in (10)–(11) reveal that increases to the buffering ratios *B*_1_ and *B_x_* relax the upper bound for integral feedback gain and thus improve stability; this suggests that non-rapid buffering of both *z*_1_ and *x*_2_ provide an effective route to suppress oscillations. As shown in SI2.2-2.3, this stabilisation effect is stronger for *b_wz_* > *γ_p_* in Topology 1 and *b_wx_* < *γ_p_* in Topology 3.

### C. Rapid Buffering with Degradation

So far we have assumed that the buffers are not subject to degradation. To establish the impact of buffer degradation on stability, here we show that buffer degradation in Topology 3 can suppress oscillations and preserve perfect adaptation; this new topology is shown in Figure 3A. Under the same assumptions as condition (8) (rapid buffering and strong binding rate constant *η*), the stability conditions are (see SI2.1):

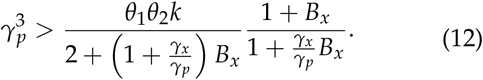

**FIG. 3.**
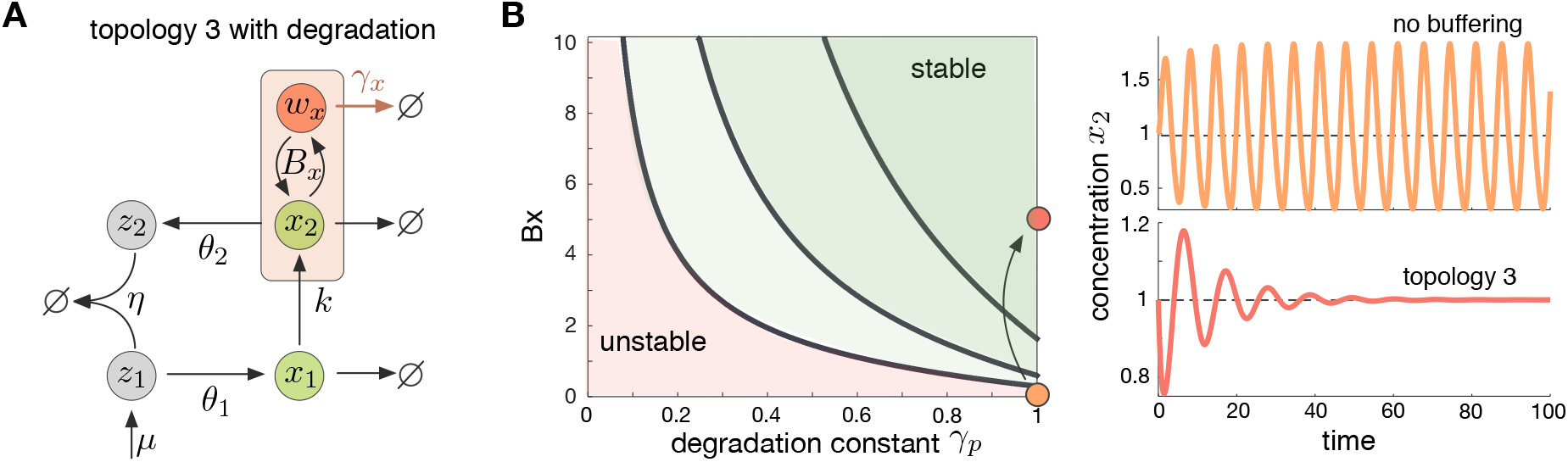
Stabilizing effect of topology 3 with degradation. **(A)** We revisit topology 3 with degradation of the buffer with degradation rate constant *γ_x_*. **(B)** Stability diagram for increasing values of the degradation rate constant *γ_x_* = {0.5,1,2}; the stability boundary corresponds to the condition in (12). Time courses are simulations of the output species (*x*_2_) for two representative cases. Parameter values are *μ* = 1, *θ*_1_ = 2, *θ*_2_ = 1, *γ_p_* = 1, *η* = 100 and *k* = 1. For the case of buffering *B_x_* = 20 and *γ_x_* = 1.

Condition (12) reduces to the one in (8) if *γ_x_* = 0 and there is no buffering of *z*_1_, i.e. *B*_1_ = 0.

As shown by the stability diagram in Figure 3B, topology 3 with degradation provides an effective solution to stabilise the closed-loop. The condition in (12) suggests that the ratio *γ_x_*/ *γ_p_* has a key role in stability. Buffers with shorter half-lives (larger *γ_x_*) tend to almost completely remove the instability, even for low buffer equilibrium ratios *B_x_*. As we show in SI2.4, under the condition *γ_x_/γ_p_* > 1/3, increases in *B_x_* tend to stabilise the closed-loop. This includes the important case where both *x*_2_ and its buffer are degraded at the same rate, i.e. *γ_x_* = *γ_p_*. For large values of *B_x_*, the stabilisation effect is even stronger and becomes independent of the half-life of *w_x_*. We found a similar stabilisation effect in systems with *x*_1_ buffering that include degradation of the buffer (see SI2.5).

## IV. ACHIEVING NEAR PERFECT ADAPTATION IN FAST GROWTH

It is well-known that dilution by cell growth can disrupt perfect adaptation in the original antithetic control system^1^. Thus here we explore the impact of dilution in the proposed topologies with molecular buffering. To study the effect of dilution, we modify (9) to include dilution terms for the control species *z*_1_ and *z*_2_, as well as the buffer for *z*_1_:

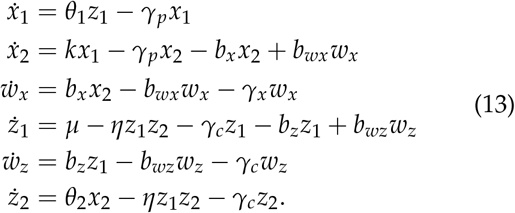

where *γ_c_* represents the dilution rate constant of the control species *z*_1_, *z*_2_ and buffer species at *z*_1_. We assume that dilution of *x*_1_ and *x*_2_ and the buffer at *x*_2_ can be lumped into their first-order degradation rates. As in the previous section, the model (13) can be further simplified for the rapid buffering case (see SI3.1).

### A. Topology 3 with dilution

We found that buffering at *x*_2_ can reduce steady state error by enabling an increase to the feedback gain without causing oscillations. For the case of dilution with a single buffer at *x*_2_, we set *b_z_* = *b_wz_* = 0 in (13). The resulting steady state is (see SI3.1):

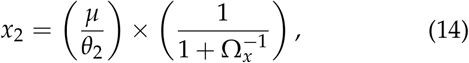

where

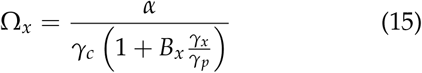

and 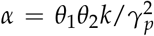. The second term on the right hand side of (14) is always smaller than unity. Therefore the steady state of the output is *x*_2_ < *μ*/*θ*_2_ and the system loses perfect adaptation. Moreover, the deviation of the steady state of *x*_2_ from the reference point *μ* /*θ*_2_ is (see SI3.1):

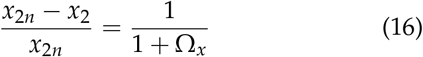

where *x*_2*n*_ = *μ*/*θ*_2_ is the reference input. Increases to *B_x_, γ_c_* or *γ_x_* in (16) thus amplify the steady state error, while increases to the feedback strength *kθ*_1_*θ*_2_ brings the system closer to perfect adaptation.

We first obtained conditions for stability in Topology 3 with dilution and non-rapid buffering (see SI3.2). Setting *γ_x_* = *γ_c_* in (13) we get:

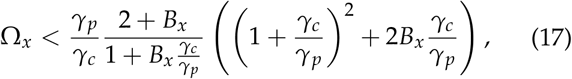

where for simplicity we have set *b_wx_* = *γ_p_* – *γ_c_*. The stability conditions for general choices of b_w_ can be found in SI3.2.

Taken together, the relations in (16)–(17) define an upper bound for the best possible steady state error. Specifically, in (16) we see that stronger feedback gain can increase Ω_*x*_ and so reduce the steady state error. Buffering of *x*_2_ tends to stabilise the oscillations and, at the same time, allows the steady state error to be reduced by stronger feedback gain, without the risk of instability observed in the original antithetic mechanism. This phenomenon is illustrated in Figure 4A, which shows the stability condition (17).

**FIG. 4.**
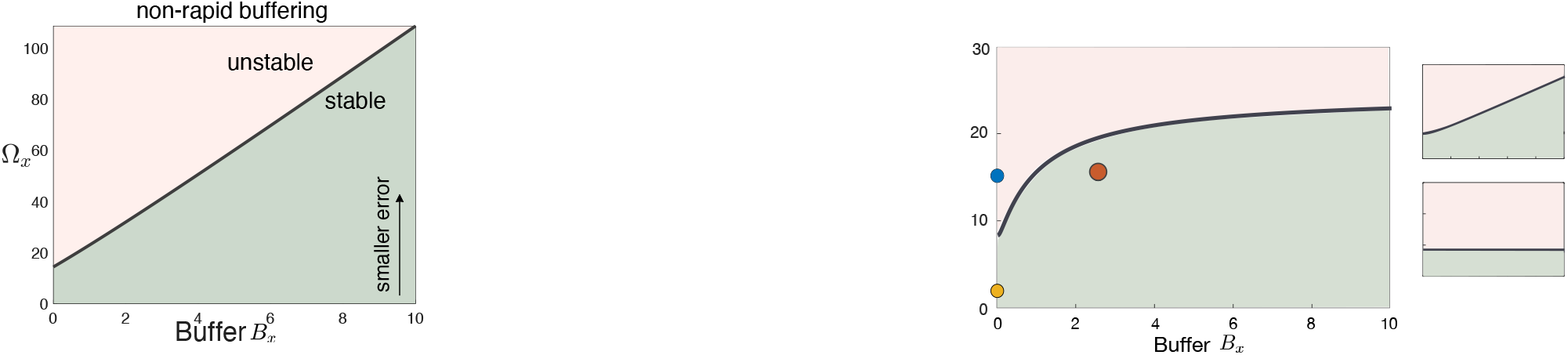
Adaptation in Topology 3 with dilution. **(A)** Stability condition in (17) for non-rapid buffering with rate constants *γ_c_/γ_p_* = 0.2. **(B)** Simulation with varying feedback gain *θ1* = {2,15,400} (from top) and buffer constant (*B_x_* = {0,2.5}). In all simulations the model parameters are *k* = 1, *θ*_2_ = 1, and *γ_p_* = 1, *γ_c_* = 1 and *γ_x_* = 10. **(C)** Stability condition in (18) for rapid buffering with rate constants *γ_c_* = 1 and *γ_p_* = 1.

In the case of rapid buffering, the condition for stability becomes (see SI3.1):

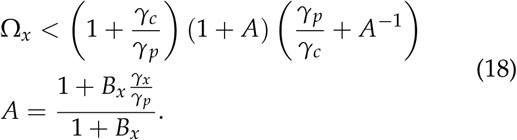

As in the non-rapid case, the relations in (16) and (18) define an upper bound for the best possible steady state error. This phenomenon is illustrated in Figure 4C, which shows the stability condition (18). Notably, we observe that increasing *B_x_* improves stability only in regions for low and high values of *γ_x_*, and not intermediate values. Figure 4B shows simulations of the stabilising effect of molecular buffering for the case of *γ_x_* = 10, which enables a decrease of steady state error by means of stronger feedback gain. We also found that topology 3 improves stability even without buffer degradation (*γ_x_* = 0), which can be observed in Figure 4B. This improvement differs from the case when there is no dilution in (8).

### B. Topology 1 with dilution

We found that non-rapid buffering at *z*_1_ can similarly reduce steady state error via increases to the feedback gain. To examine this result in detail, we set *B_x_* = 0 in (13) and compute the resulting steady state (see SI3.4):

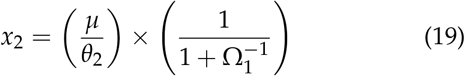

where

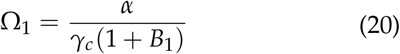

and 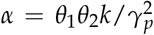. As in the previous case, the steady state satisfies *x*_2_ < *μ*/*θ*_2_ and thus the system loses perfect adaptation (see Figure 5). Moreover, in this case the steady state error is (see SI3.4):

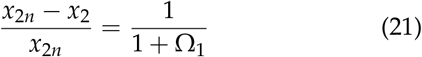

where *x*_2*n*_ = *μ*/*θ*_2_ is the reference input. Increasing *B*_1_ or *γ_c_* in (21) increases the steady state error, while increasing the feedback strength *kθ*_1_*θ*_2_ brings the system closer to perfect adaptation. If for simplicity we assume that *b_wz_* = *γ_p_* – *γ_c_*, the condition for stability is (see SI3.5):

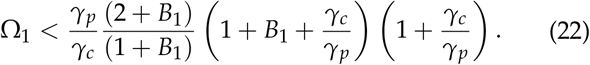

**FIG. 5.**
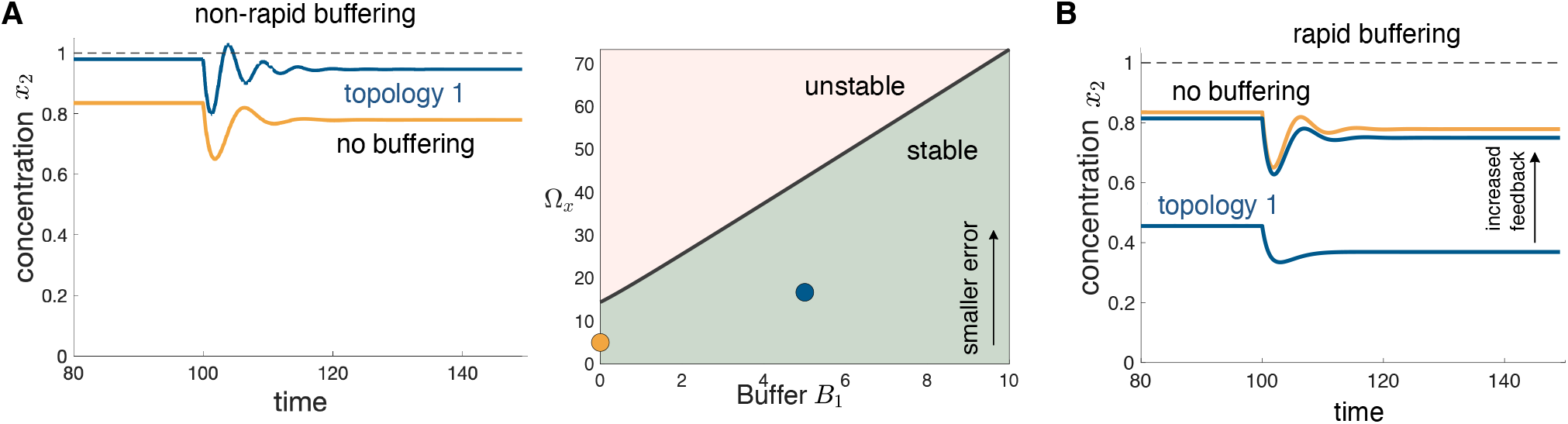
Adaptation in Topology 1 with dilution. **(A)** Non-rapid buffering with stronger feedback reduces the adaptation error. (Left) In simulations the model parameters are *θ*_2_ = 1, *γ_p_* = 1, *η* = 100, *μ* = 1, *γ_c_* = 0.2, and *k* = 1 changes to *k* = 0.7 at *t* = 100. (Right) The stability condition in (22) for non-rapid buffering with dilution rate constant in the ratio *γ_c_/γ_p_* = 0.2. **(B)** Rapid buffering increases the adaptation error while rapid buffering with stronger feedback is identical to no buffering with weaker feedback. Parameter values for the non-rapid case are *θ*_1_ = {1,15}, *B*_1_ = {0,5} and *b*_*w*1_ = {0,0.8}; parameters for the rapid case are *θ*_1_ = {1,5} and *B*_1_ = {0,5}.

The general case for other choices of *b_w_* can be found in SI3.5. Non-rapid buffering thus enables a reduction of the steady state error via increased feedback gain without unwanted oscillations. This phenomenon is illustrated in Figure 5A, which shows simulations and the stability condition (22) that describes the upper bound on the steady state error. In contrast, rapid buffering does not improve the stability condition and so does not enable a decrease in the adaptation error (see SI3.4); we have illustrated this phenomenon in Figure 5B via numerical simulations.

While the benefit of buffering described here are for a simple network and a steady state error (adaptation) tradeoff, control theory can be use to mathematically quantify this benefit for more general networks and more general tradeoffs involving both steady state and temporal dynamics (see SI4.3).

## V. CASE STUDIES

### A. Model for biofuel production

To illustrate the potential of the proposed control topologies, here we employ an existing model for biofuel production that incorporates antithetic control^6^ (see also^12^), shown in Figure 6A. The synthetic system produces biofuel from sugars through a metabolic pathway. The biofuel product can be toxic to the cell and so efflux pump proteins are expressed to remove the toxic metabolic product. However, at large concentrations the efflux protein pump can also be toxic. A feedback mechanism can help robustly regulate these two competing toxic products. The antithetic feedback mechanism senses the biofuel concentration to control the expression of efflux pump protein. An increase in the pump protein then reduces the biofuel concentration, completing the loop. Stability is known to be a major issue for the system, as it susceptible to oscillation for large *η*, which is the typical design case^6^.

**FIG. 6.**
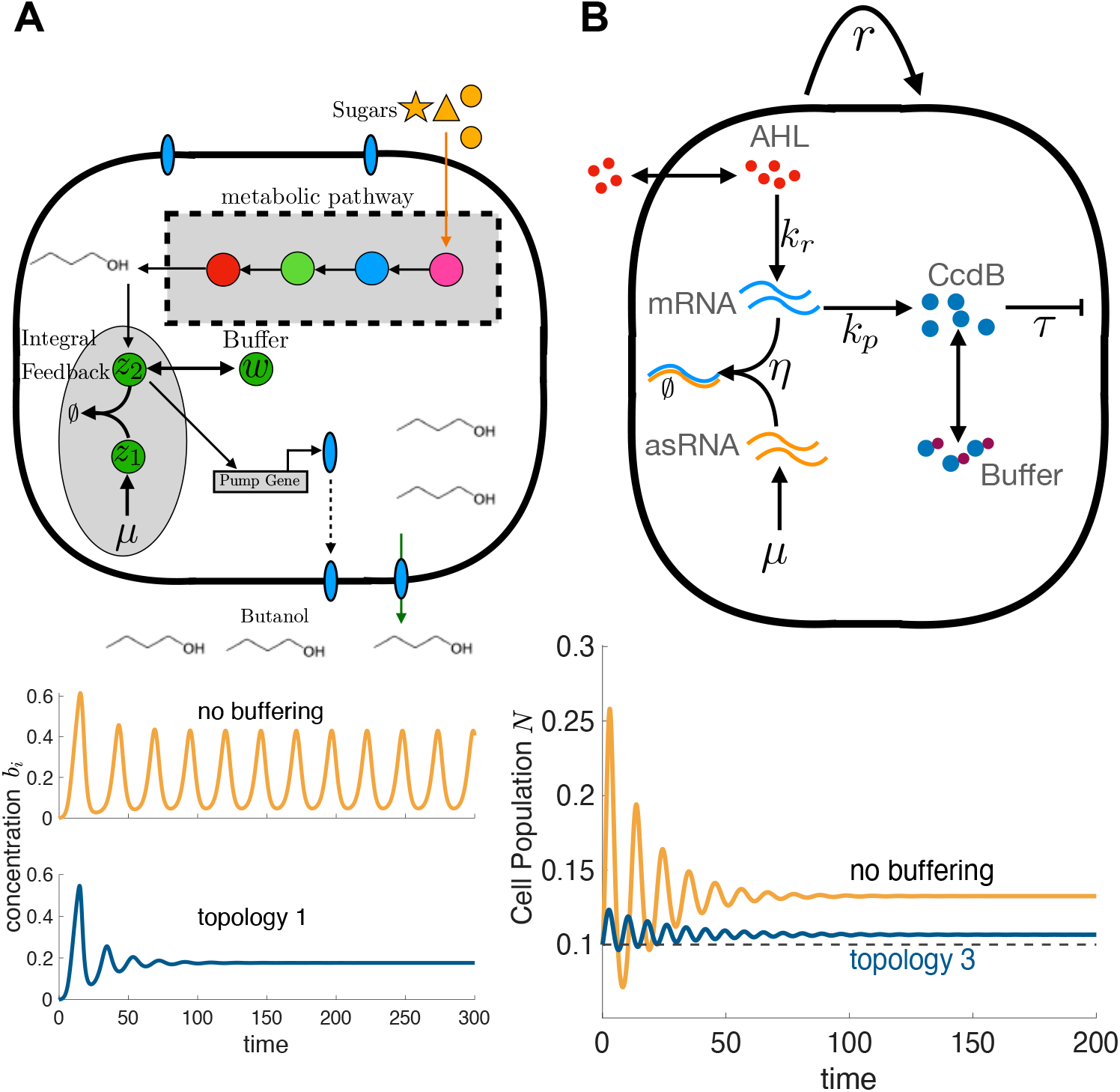
Case studies of molecular buffering coupled with antithetic control showing improved stabilisation and adaptation. **(A)** Biofuel production system adapted from^6^ to include buffering of species *z*_2_. Simulations show the stabilising effect of buffering. Model parameters for simulations are *α_n_* = 0.66, *δ_n_* = 0.5, *γ_p_* = 0.14, *α_b_* = 0.1, *δ_b_* = 0.5, *β_p_* = 0.66, *V* = 1, *η* = 100, *μ* = 0.1762, *θ* = 1, *I* = 1, *r_p_* = 0,1.25, *r_w_* = 0,0.25, *k* = 0.5,3. *k* is increased with buffering to compensate for the reduced integral feedback gain from buffering **(B)** Synthetic growth control circuit^27^ adapted to include buffering of CcdB. Simulations show the ability of buffering to decrease the steady state error via stronger feedback without oscillations. Model parameters for simulations with and without buffering are *γ_p_* = 3, *r* = 1, *τ* = 4 × 10^−3^, *k_R_* = 10^−1^, *G* = 10^−6^, *η* = 20, *γ_r_* = 0.1, *μ* = 10, *N_m_* = 10^9^, *γ_w_* = 10^−2^, *k_p_* = {20,200}, *b_c_* = {0,7} and *b_w_* = {0,7 × 10^−2^}.

The extended model of the biofuel circuit with antithetic feedback and the addition of a protein buffer is:

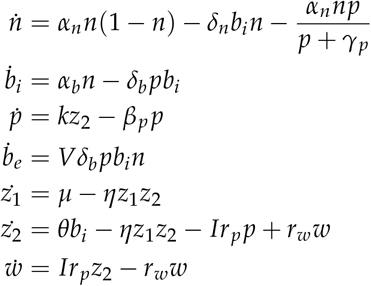

where *n* is the normalized cell density, which is assumed to follow logistic growth with additional death rates due to toxicity of intracellular biofuel concentration *b_i_* and efflux protein pump *p*. The variables *z*_1_ and *z*_2_ are the controller species, while the production of the protein pump *p* is assumed to be proportional to controller species *z*_2_. The variable *w* is the buffering species which buffers *z*_2_ through a reversible reaction via chemical species *I* that inhibits *z*_2_ sequestering when bound to *z*_2_. The variable *b_e_* is the extracellular concentration of biofuel.

Buffering of *z*_2_ can be seen to stabilise the process in the simulations of the model (see Figure 6A). These simulations show that the oscillations, which occur when *η* is too large, quickly settle to the steady state when buffering is introduced. This stabilising effect is equivalent to the impact of buffering in Topology 1 in Section III. Buffering removes the stability limit on *η* and so a large antithetic binding rate *η* is possible without oscillations. This example thus illustrates that the stabilising effect of buffering also occurs in more complex systems than for the simple case presented above.

### B. Model for growth control

For the synthetic growth control case study, we use an existing model of the synthetic growth control circuit, which includes the new addition of buffering^27^ (see Figure 6B). The variable *N* represents the population size and is assumed to follow logistic growth, with an additional death rate due to toxicity that is proportional to the concentration of *CcdB* per cell. *Ccdb* is a protein that is toxic to the cell. *mRNA* is messenger RNA while asRNA is a short antisense RNA that has a complementary sequence to the mRNA, which enables sequestration between the two. *mRNA* and *asRNA* form the antithetic integral controller. The transcription of mRNA is induced by a quorum-sensing ligand. The term *G_a_* represents the gain between *N* and mRNA induction resulting from the quorum-sensing molecule AHL. *W* represents a buffer of Ccdb, which consists of an inactivated form of Ccdb that can reversibly bind to an inhibitor molecule *I*. The adapted model of the genetic circuit with the new addition of a protein buffer is

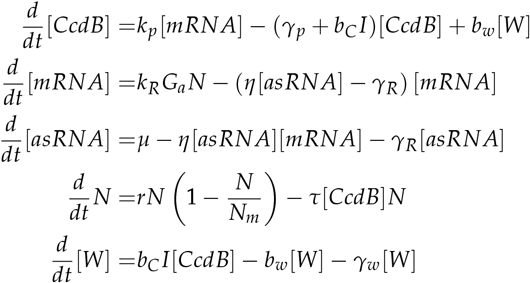

where [·] represents intracellular concentrations for each species and the last line indicating the rate of change of *W* is new to the model.

The buffering of *CcdB* as shown above is equivalent to *x*_1_ buffering in the model (6) and Figure 2, as *N* is the output and equivalent to *x*_2_. Buffering at *x*_1_ provides a similar benefit as buffering at *x*_2_ and so can also enable near-perfect adaptation.

Buffering of *CcdB* in conjuction with increased feedback gain can be shown to reduce steady state error in the simulations in Figure 6. Increased feedback gain is implemented in these simulations by increasing the translation rate of *CcdB*.

## VI. DISCUSSION

Perfect adaptation has been subject of intense study in the synthetic biology community. Although perfectly adapting systems are ubiquituous in nature, their implementation has proven particularly elusive. The antithetic control motif, first discovered by Briat et al^4^ and implemented by Aoki et al^1^, provides a new molecular mechanism to build perfect adaptation into a wide range of synthetic gene circuits. A number of works have sought to find alternative circuits that provide adaptation properties similar to antithetic control. For example, several authors have shown that ultrasensitive feedback can display some of the features of perfect adaptation^25,28^, and the idea was recently extended in great detail for synthetic gene circuits^30^. Other works have sought to devise molecular implementations of Proportional-Integral-Derivative control^9^, as this is a widely adopted strategy for perfect adaptation in engineered control systems.

Here we have addressed caveats of the original antithetic control system with an extended architecture that has improved stability properties. The proposed circuit combines an antithetic motif with a molecular buffering mechanism. Molecular buffering is widely conserved in natural systems, and common examples include the ATP buffering by creatine phosphate, pH buffering and calcium buffering. In all these examples, a molecular buffer sequesters a target molecule into an inactive form, resulting in a system with improved ability to mitigate fast perturbations. In the case of antithetic control, the addition of buffering results in the stabilisation of unwanted oscillations and, moreover, provides near-perfect adaptation even in rapid growth conditions where the performance of antithetic control is known to be particularly poor.

After detailed examination of mathematical models for various circuit architectures (Fig. 2) with rapid and non-rapid buffers, we found two candidate systems with improved stability properties, either by buffering species of the antithetic motif itself, or by buffering a target species to be controlled. The first circuit, called Topology 1 in Fig. 2, under non-rapid buffering provides stability over a larger range of parameters values than classic antithetic control and can generally stabilise unwanted oscillations. Moreover, Topology 1 requires buffering of a molecular species of the antithetic motif itself, and therefore it provides a promising strategy to stabilise variables that are not easily buffered directly, such as population size or metabolite species as illustrated by the example in Fig 6A.

The second circuit, termed Topology 3 in Fig. 2, requires buffering the molecular output of the process to be controlled. We found that when *x*_2_ buffering is either non-rapid or rapid and coupled with degradation, it mitigates oscillations in fast growth regimes. Interestingly, there is a similar effect when applied to intermediate species instead of the output species in the controlled process, such as *x*_1_ in the original circuit shown in Fig. 2 or CcdB in the growth control case study in Fig. 6. Buffering an intermediate species also provides additional design flexibility when the output species cannot be easily buffered.

Although our results indicate that both buffering and degradation can act as stabilisers of antithetic control, designs with increased buffering and reduced degradation can provide substantial benefits. Increased degradation requires a higher production rate to achieve a particular steady state concentration. If the degradation mechanism requires expression of heterologous proteins, an increase in their production rate imposes a heavier genetic burden on the host cell^8,26^. Moreover, in applications, protein production rates are subject to upper limits depending on the genetic machinery of the host^36^, and so increasing degradation can also limit the maximum set point concentration. Tuning of degradation can also be difficult to implement given the limited number of degradation tags available.

The effect of buffering on adaptation is strikingly similar to a strategy employed in industrial process control, where buffer tanks are employed to regulate and smooth out the impact of disturbances^13^. In our case, the specific implementation of the molecular buffers is a subject of future study, as this will largely depend on the type of biomolecular process to be controlled (see^18^ for quantification of different buffering reaction forms). For example, buffers for gene expression may require gene products to be sequestered, which can be achieved through several mechanisms such as reversible protein-protein binding^24^, phosphorylation^19^, small molecule inhibitors^11^, or DNA decoy sites^34^. In metabolism and signalling systems, ubiquitous examples are the interconversion between a target species and a buffer (e. g. reversible catalysis between ATP and creatine phosphate^17,32^) or sequestering by dedicated proteins (e. g. Ca^2+^ or H^+^ ions^17,21,32^).

Our main goal in this paper was to show that molecular buffering can improve perfect adaptation in the antithetic control motif. Since buffering is known to stabilise a much wider range of molecular networks^16^, it also has the potential to improve other circuits implementing perfect adaptation, e. g. those that rely on ultrasensitive behaviour^30^. Further, its properties may be used to instead improve transient response or disturbance rejection of a circuit, rather than only adaptation^16,17^. A key future step is the study of design rules for buffering and antithetic feedback. A useful starting point in this direction are the rules from lead-lag and PID controller tuning from control engineering^16^ (see SI4.2). Unlike technological controllers, however, the stabilisation properties of buffering are tied to its synergy with feedback regulation^16,17^ and the location of the buffer in a network is important, which introduces new challenges for controller design.

For simplicity, here we have focused exclusively on deterministic dynamics, but the analysis of stochastic effects emerging from the interplay between molecular buffering and antithetic control are particularly attractive, as it is known that buffering does not amplify stochastic fluctuations^17^ yet some phenomena are known to emerge only in the presence of molecular noise^4,31,33^.

As synthetic gene circuits grow in size and complexity, there is a growing need for mechanisms that can enhance their robustness in a range of operational conditions. In the longer term, this will require the availability of a catalogue of gene circuits that can produce perfect adaptation in response to perturbations. In this work we have presented one such architecture, and thus laid theoretical groundwork for the discovery of biomolecular systems with improved functionality.

## METHODS

All mathematical models are based on systems of ordinary differential equations. The stability conditions in Eqs. (8), (12), (18) and (22) were obtained using frequency domain transformations (Laplace and Fourier) of the linearised models, along with detailed examination of the magnitude and phase equations of the resulting characteristic polynomials for the closed-loop systems^27^. Simulations were carried out using standard ODE solvers in MATLAB. All calculations and model descriptions can be found in the Supplementary Material.

## AUTHOR CONTRIBUTIONS

Conceptualization, E.J.H.; Methodology, E.J.H.; Formal Analysis: E.J.H., and D.A.O.; Investigation, E.J.H., and D.A.O.; Writing - Original Draft, E.J.H.; Writing - Review & Editing, E.J.H., and D.A.O.; Funding Acquisition, E.J.H.; Supervision, E.J.H., and D.A.O.

## ACKNOWLEDGEMENTS

The authors would like to thank Morgan Kelly for her assistance. EJH was would like to thank the contribution of Judith and David Coffey.

The authors declare that there are no competing interests.

## Supplementary material

### SI1. BUFFERING BACKGROUND

In this section, we provide a background on buffering^4^, including methods for analysing models with buffering. We start with the simple model in the main section of a single regulated species that is being buffered, such that

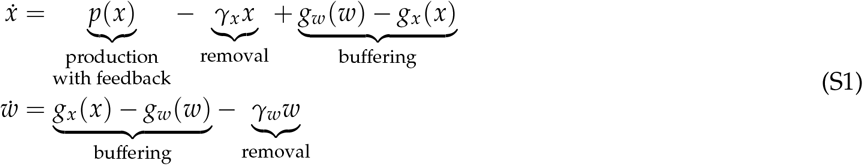

where *x* is the output species being regulated, *w* is the buffering species, *p* is the production rate of *x, γ_x_* is the removal kinetic rate of *x, g_w_* is the forward buffering reaction rate and *g_x_* is the reverse buffering reaction rate. Incorporation of feedback is represented by the *x* dependence of production. The steady state occurs when production matches degradation 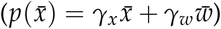 and the buffer is at steady state 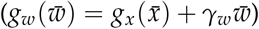. To analyse (S1), we reduce the two state model to one state by assuming that the buffering reactions rapidly reach equilibrium (see^4^ for more mathematically rigorous derivation). To carry this out, we first linearise (S1), which results in

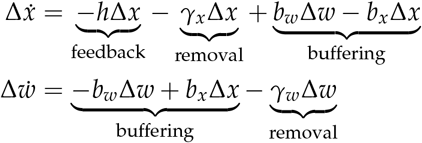

where 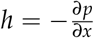 is the linearised feedback gain, and 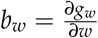 and 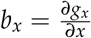 the linearised kinetic rates for the forward and reverse buffering reaction.

We introduce the variable Δ*x_T_* = Δ*w* + Δ*x* to carry out a quasi-steady state approximation^2,4^, where we have the transformed model

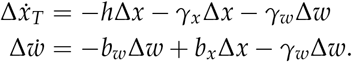

We assume that the buffering species Δ*w* rapidly reaches quasi-steady state due to large *b_w_* and thus assume 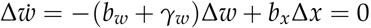, where *x_T_* is a ‘slow’ variable as it is not a direct function of the buffering reactions. The quasi-steady state can be rewritten

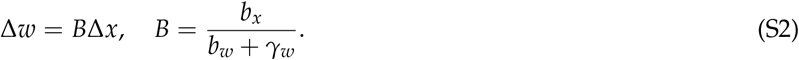

From this quasi-steady state, we have Δ*x_T_* = (1 + *B*)Δ*x* and Δ*ẋ_T_* = (1 + *B*)Δ*ẋ*, resulting in

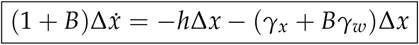

which is a reduced one state model, where the second state can be determined from Δ*w* = *B*Δ*x*.

In technology, integral feedback is often paired with proportional and derivative feedback (PID control)^1^. In biology, rapid buffering without degradation is equivalent to negative derivative feedback and rapid buffering with degradation is equivalent to PD feedback with degradation. These equivalences can be observed in^4^

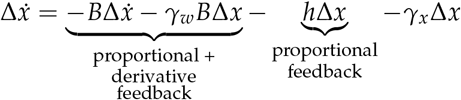

where the buffer equilibrium ratio *B* corresponds to the derivative feedback ‘gain’.

To study the effect of buffering on stability, we can also modify the model in (S1) to include a delay of the production feedback term

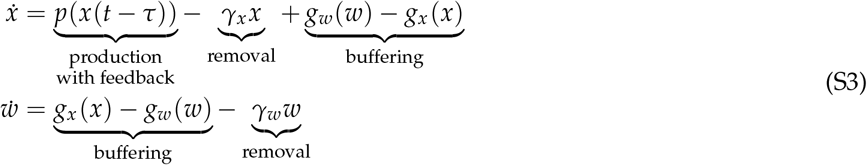

where *p*(*x*(*t* – *τ*)) represents the production feedback with a delay of time *τ*. The reduced model is

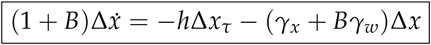

where Δ*x_τ_* = Δ*x*(*t* – *τ*).

It can be observed in Figure S2 that buffering can stabilise the oscillations that result from feedback delay. This stabilisation is a result of (a) the buffering reactions acting directly on the target molecular species (i.e. not being subject to delays) and (b) buffering having a ‘derivative feedback’ effect - known for its stabilising effects in control engineering^1^.

**FIG. S1.**
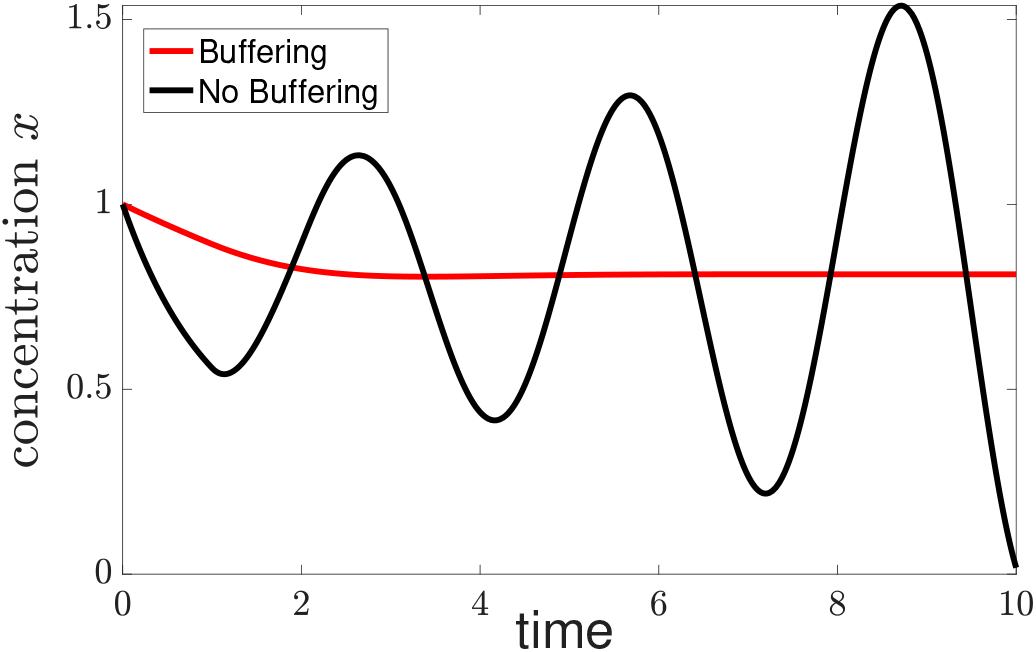
The parameters are *B_x_* = 0 and 5, *τ* = 1 (delay), = 1, *γ_w_* = 0 and *h* = 2.7.

### SI2. STABILITY OF ANTITHETIC INTEGRAL FEEDBACK WITH BUFFERING

#### SI2.1. Stability of Antithetic Integral Feedback with Rapid Buffering: All Species

In this section, we analyse the stabilising effect of buffering (without degradation) on antithetic integral feedback. We base our studies on a simple model involving the antithetic integral feedback (without buffering)^5^. Consider

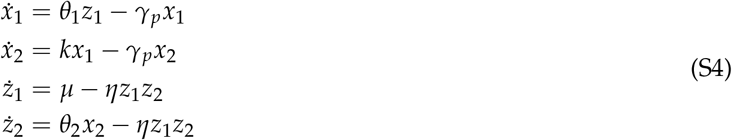

where *x*_2_ is the output concentration being controlled, *x*_1_ is another concentration in the process being controlled, and *z*_1_ and *z*_2_ represent the molecular species involved in the perfect adaptation mechanism.

We next introduce buffering to (S4). We show how the model reduction method described in SI1 can be used to simplify the model for one case. We then use the same method for all buffers.

As a first case, we introduce buffering at the controlled variable *x*_2_ and simplify the model by assuming that the buffering reactions rapidly reach equilibrium. With buffering at *x*_2_, we have

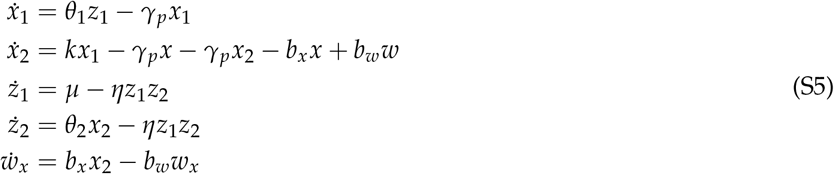

where *w_x_* is the buffering species at *x*_2_ and *b_x_, b_w_* are the kinetic rate constants for the buffering reactions. Although the buffering equilibrium ratio is defined in terms of the linearised model and deviations from steady state, we can use the same notation here for the nonlinear model as the buffering reactions are linear. If the buffering reaction is at equilibrium then

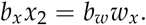

If we assume that the buffer rapidly reaches equilibrium then *w* is at quasi-steady state and so

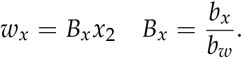

We note that although the buffer equilibrium ratio is defined in (S2) in terms deviations from steady state, we can use the buffer equilibrium ratios in the nonlinear model as the reaction rates are linear.

We set *x_T_* = *w_x_* + *x* as the slow variable and so *x_T_* = (1 + *B_x_*)*x*_2_. Thus *ẋ_T_* = (1 + *B_x_*)*ẋ*_2_ and so

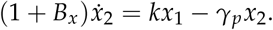

If we include buffering on *z*_1_, *z*_2_, *x*_2_ and apply a similar model reduction by assuming rapid buffering, then we have

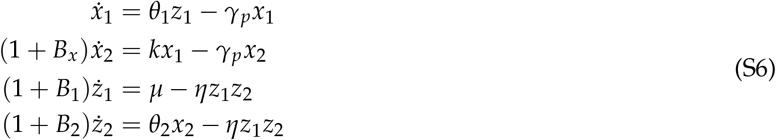

where *B*_1_, *B*_2_ are the buffer equilibrium ratios of the buffers at *z*_1_ and *z*_2_ respectively.

##### SI2.1.a. Steady State

We first determine the steady states of the system, which is useful both for determining perfect adaptation and as a prerequisite for stability analysis. Using

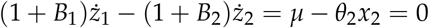

we can see that the steady state for the output is

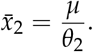

The correspondence of this steady state with perfect adaptation is discussed further in Section 2 & 3 of the paper. The species *x*_1_, *z*_1_ and *z*_2_ have the corresponding steady states

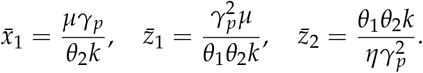

##### SI2.1.b. Linearised Model

We next determine the effect of buffering on the stability of the model. We follow the methodology used by Olsman and colleagues^5^ to study the stability of antithetic integral feedback with the addition of buffering. Linearising the system (S6) about the steady states, we have

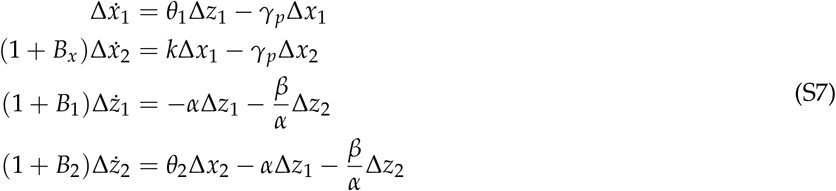

where 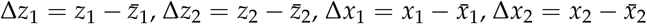 are the deviations from steady state and

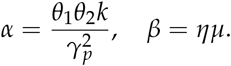

Taking the Laplace transform of (S7), we have

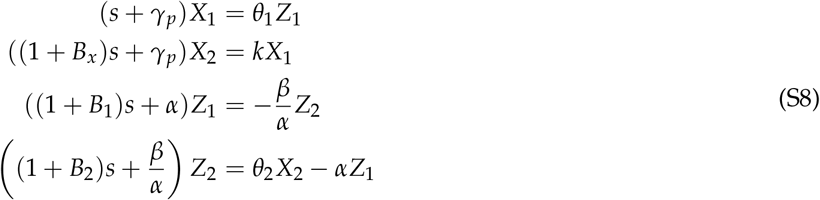

where 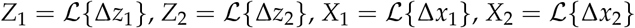 are the Laplace transforms of the time-domain concentration deviations. Substituting, we have

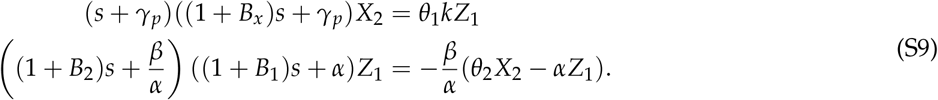

Simplifying, we have

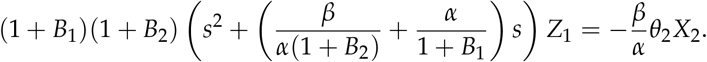

Substituting from (S9), we have

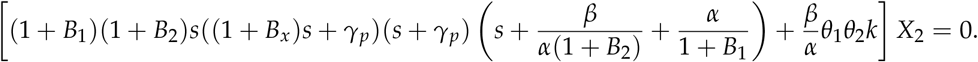

The characteristic equation used to analyse stability is thus

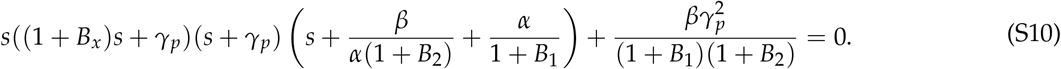

##### SI2.1.c. Characterisation of roots

Following the methodology used by Olsman and colleagues^5^, we first characterise the roots of (S10). If we substitute *s* = *γ_p_σ* then

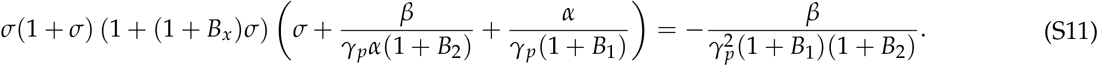

Taking the limit of strong binding for the sequestration process in antithetic integral feedback 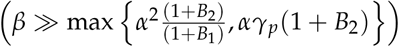 then

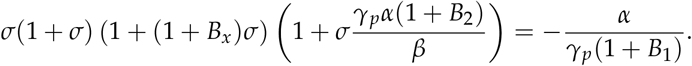

It can be observed that there is a large, real root at 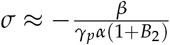. We next examine the region 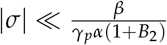, where we have

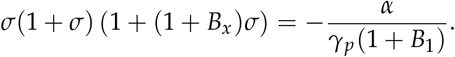

The magnitude and phase constraints are

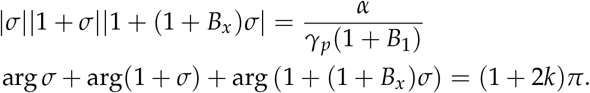

If we assume that *σ* is real and positive, then the LHS of the phase constraint is

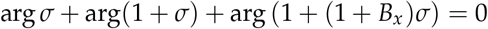

which contradicts. Thus unstable roots are not purely real.

If we assume that *σ* is real and −1/(1 + *B_x_*) < *σ* < 0 then the LHS of the phase constraint is

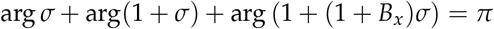

and so it is possible to have stable real roots. If we set *f* = |*σ*||1 + *σ*|| 1 + (1 + *B_x_*)*σ*| from the magnitude constraint, then there is a local maxim of *f* at

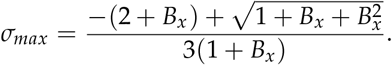

Thus there are two stable real roots between −1/(1 + *B_x_*) < *σ* < 0 if

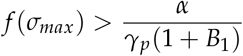

as *f*(*σ_max_*) is larger than the RHS of the magnitude constraint. There is a bifurcation at the boundary 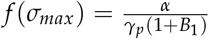 resulting in a pair of complex conjugate roots if

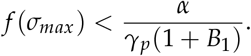

For −1 < *σ* < −1/(1 + *B_x_*) then

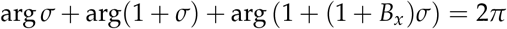

which violates the phase constraints. For *σ* < −1 then

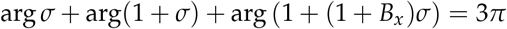

which meets the phase constraints. Thus for the stability boundary with strong binding there is a negative real root and a complex pair of roots in the region 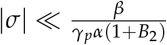, as well as one large negative root.

##### SI2.1.d. Stability Conditions

We next determine the stability boundary, where roots of the characteristic equation have zero real parts. Substituting *s* = *iωγ_p_*, we have

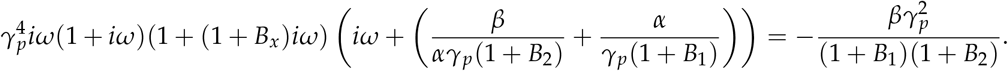

From these equations, the magnitude and phase constraint are

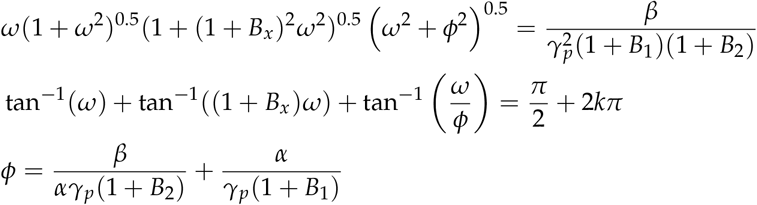

for some integer *k*. Using the strong binding assumption 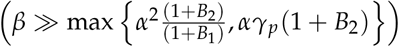 then from above 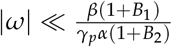 and so

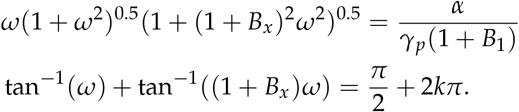

Rewriting the phase constraint, we have

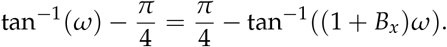

Applying tan(·) and trigonometric identities to both sides, we have

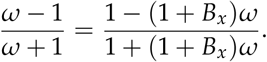

Solving, we have

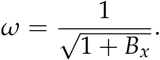

Thus the stability boundary occurs at

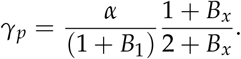

From above, we know that all roots are real and stable if *α*/(*γ_p_*(1 + *B*_1_)) is sufficiently small, and so the stability condition is

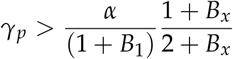

or

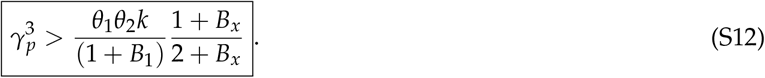

Thus increasing *B*_1_ improves stability and increasing *B*_2_ has no effect on stability. Further, increasing *B_x_* worsens stability, although this effect saturates as *B_x_* increases.

#### SI2.2. Stability of Antithetic Feedback with Non-Rapid Buffering at *z*_1_

In this section, we analyse the ability of non-rapid buffering at *z*_1_ to stabilise antithetic integral feedback. We use the model

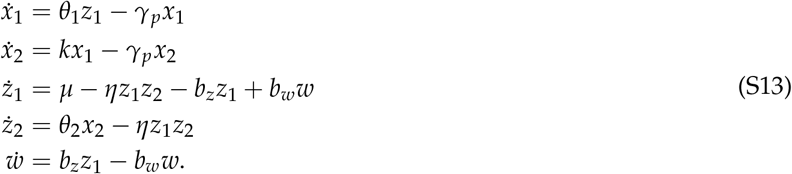

where the buffer *w* is not assumed to rapidly reach equilbrium. As a result, the model cannot be reduced in a similar manner to SI2.1. The steady state for (S13) is identical to the rapid case in SI2.1. The linearisation is

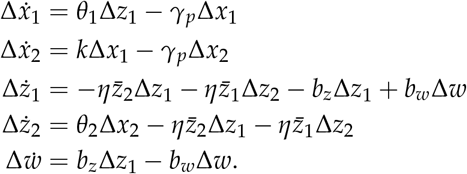

This system can be rewritten

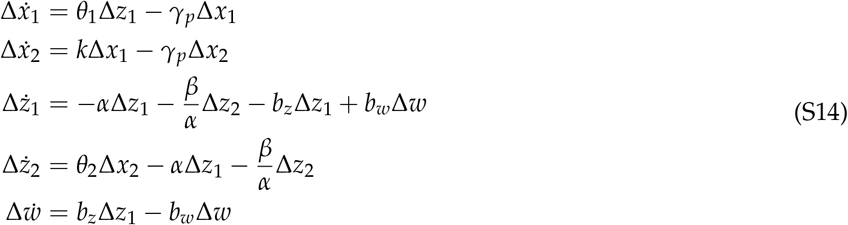

where

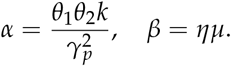

Taking the Laplace transform of 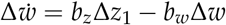, we have

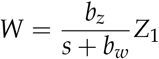

where *W* and *Z*_1_ are the Laplace transforms of *w* and *z*_1_. We have

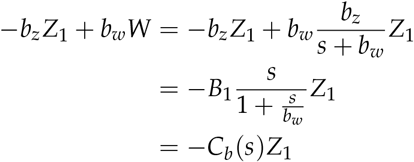

where

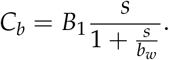

Taking the Laplace transform of (S14), we have

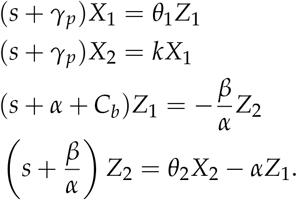

Combining, we have

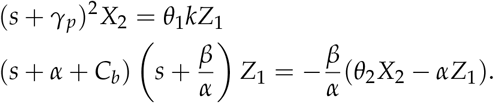

Simplifying, we have

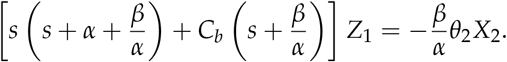

Taking the strong antithetic binding limit of the sequestration mechanism (*β* ≫ max {*α*^2^, *γ_p_α*}), we have

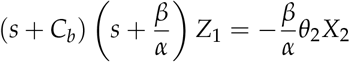

and so

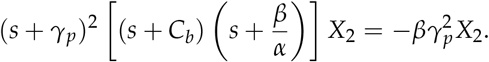

Rewriting *C_b_*, we have

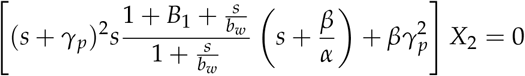

and so the characteristic equation is

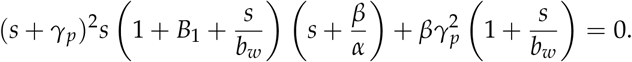

Substituting *s* = *iωγ_p_*, we have

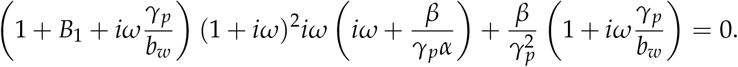

Taking the strong antithetic binding limit where 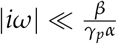, we have

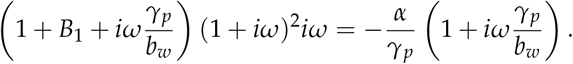

The magnitude constraint is

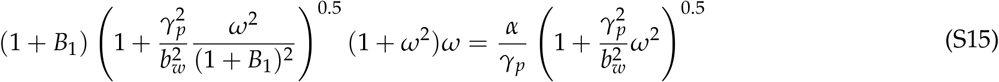

and the phase constraint is

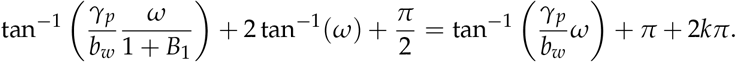

for some integer *k*. Using trigonometric identities, we have

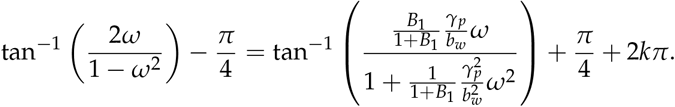

This can be simplified to

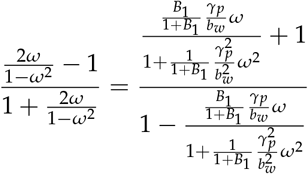

or

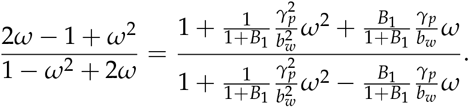

We have

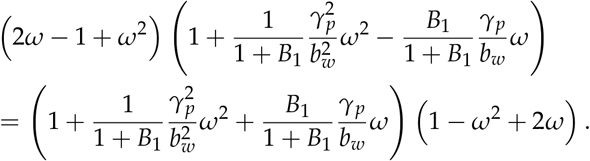

Setting

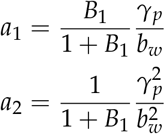

we have

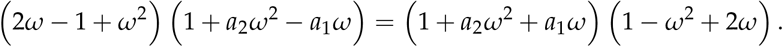

Matching coefficients, we have

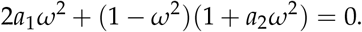

and thus

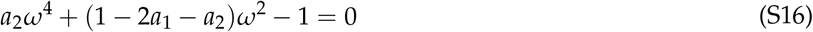

Solving for *ω*^2^, we have the stability constraint

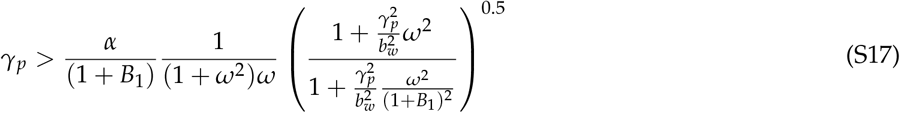

where

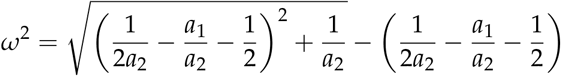

which can be rewritten as

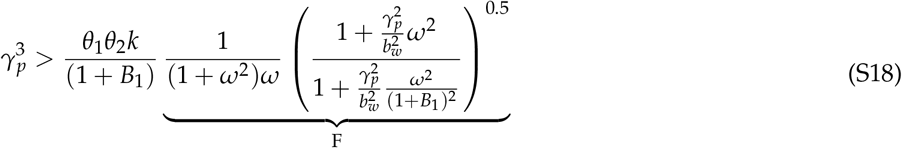

**FIG. S2.**
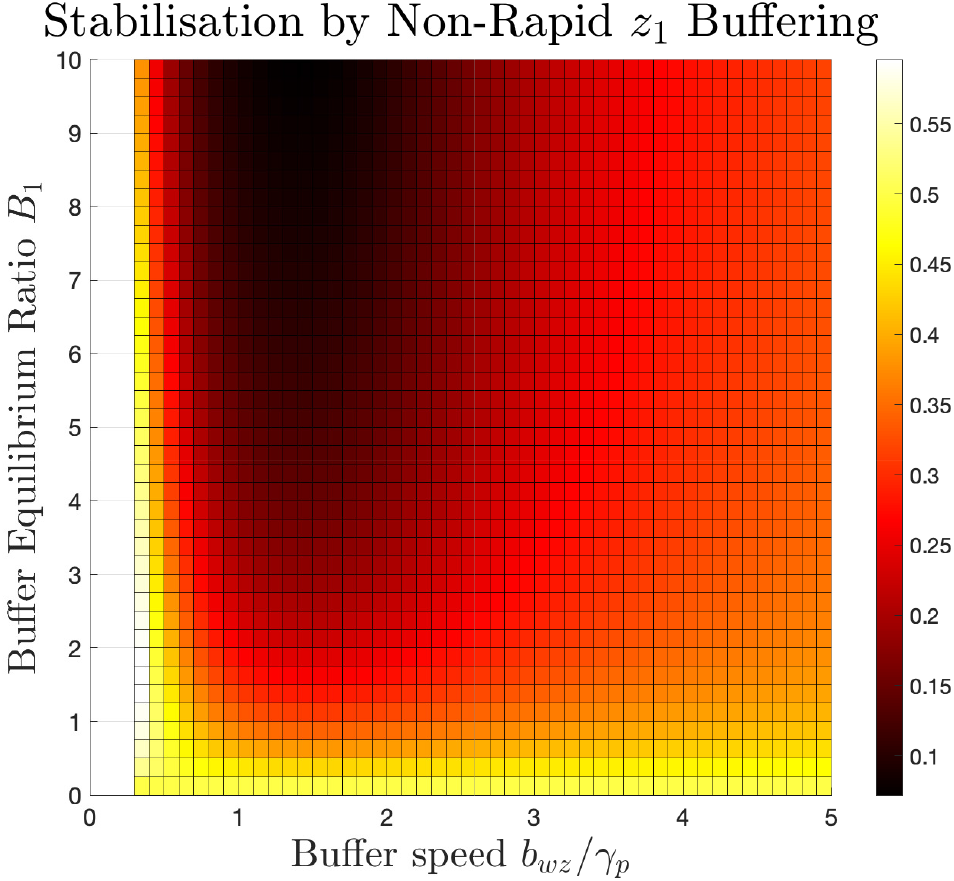
Stabilisation effect F in (S18) where smaller implies improved stability.

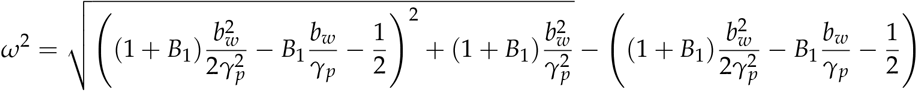

where *F*, which is the stabilisation effect independent of the integral gain, can be seen as a function of *B*_1_ and 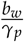 in Figure S2.

For the special case that *b_w_* = *γ_p_* then we have

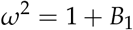

and so

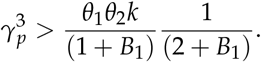

#### SI2.3. Stability of Antithetic Feedback with Non-Rapid Buffering at *x*_2_

In this section, we analyse the ability of non-rapid buffering at *x*_2_ to stabilise antithetic integral feedback. We use the model

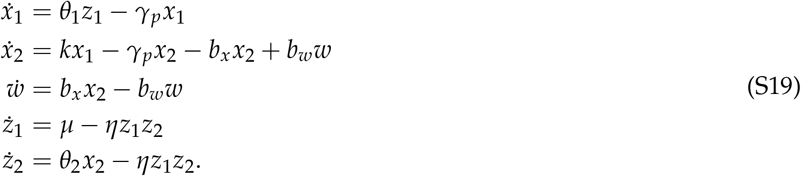

where the buffer *w* is not assumed to rapidly reach equilbrium. As a result, the model cannot be reduced in a similar manner to SI2.1. The steady state for (S19) is identical to the rapid case in SI2.1. The linearisation is

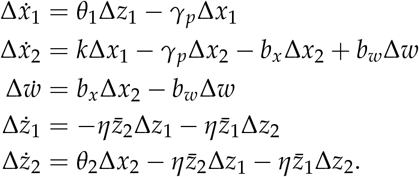

This system can be rewritten

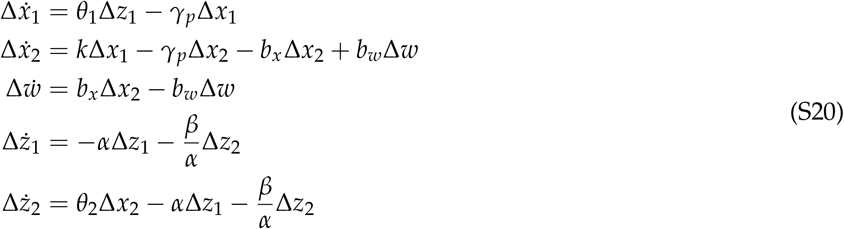

where

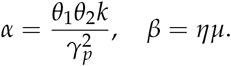

Taking the Laplace transform of 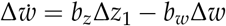, we have

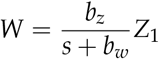

where *W* and *Z*_1_ are the Laplace transforms of *w* and *z*_1_. We have

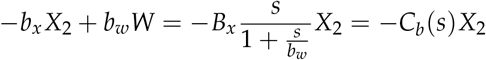

where

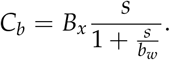

Taking the Laplace transform of (S20), we have

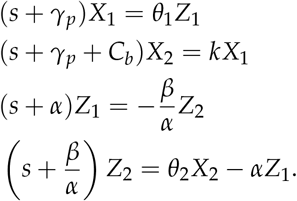

Combining, we have

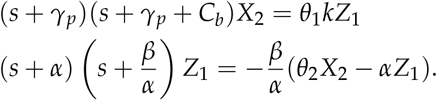

Simplifying, we have

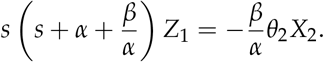

Taking the strong antithetic binding limit of the sequestration mechanism (*β* ≫ max {*α*^2^, *γ_p_α*}), we have

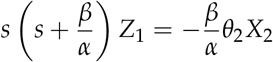

and so

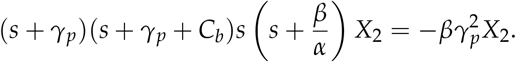

Rewriting *C_b_*, we have

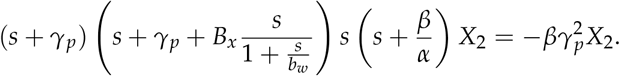

and so the characteristic equation is

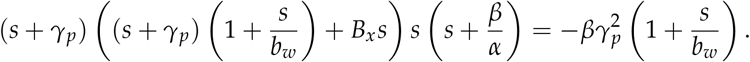

Substituting *s* = *iωγ_p_*, we have

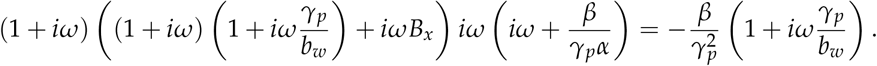

Taking the strong antithetic binding limit where 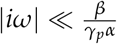, we have

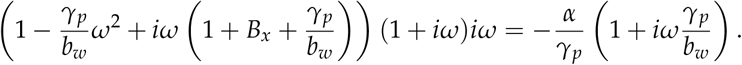

The magnitude constraint is

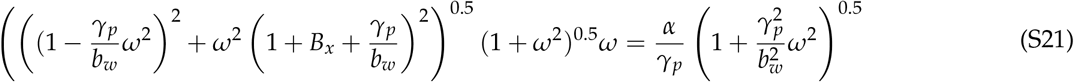

and the phase constraint is

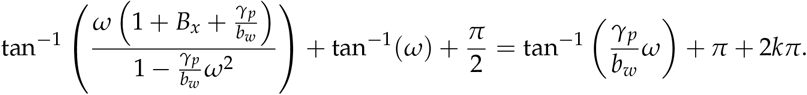

for some integer *k*. Using trigonometric identities and rearranging, we have

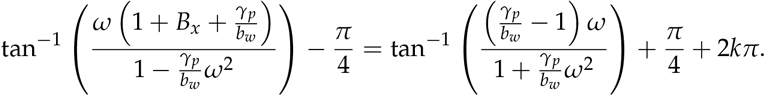

Taking the tangent of both sides, we have

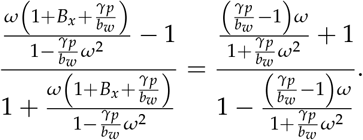

Multiplying out fractions, we have

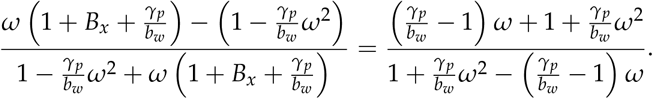

Simplifying, we have

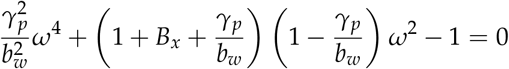

**FIG. S3.**
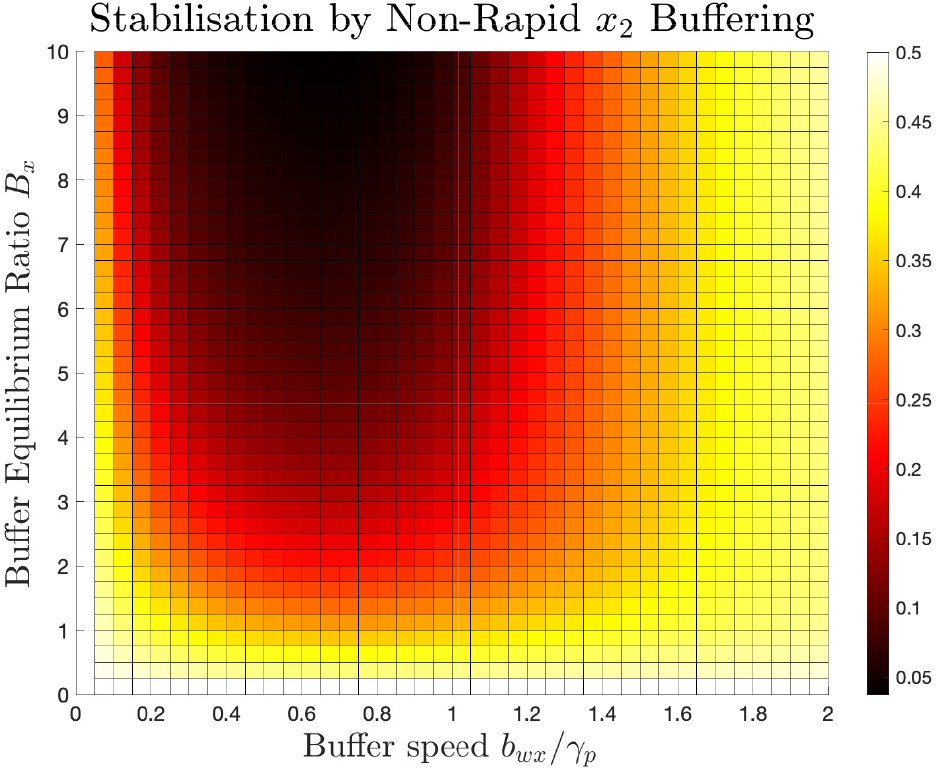
Stabilisation effect F in (S22) where smaller implies improved stability.

Solving for *ω*^2^ and using the same approach as in previous sections, the magnitude constraint leads to the stability

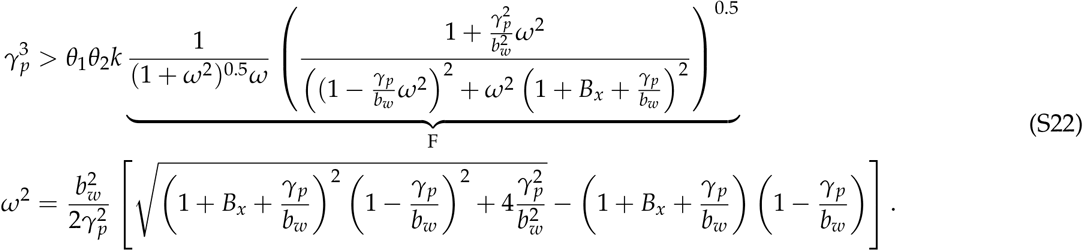

where *F*, which is the stabilisation effect independent of the integral gain, can be seen as a function of *B_x_* and 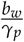 in Figure S3.

For the special case where we set *b_w_* = *γ_p_* then we have *ω*^2^ = 1 and so the stability constraint is

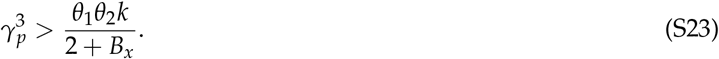

#### SI2.4. Stability of Antithetic Feedback: Rapid *x*_2_ Buffering with Degradation

In this section, we analyse the stabilising effect of buffering at the output species *x*_2_ on antithetic integral feedback, where the buffering can be degraded. Consider the model (S5) with buffering at *x*_2_ that is degraded

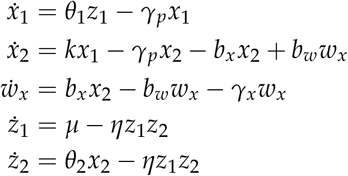

where *w_x_* is the buffering species of *x*_2_, and *b_x_, b_w_* are the kinetic rates for the buffering reactions. If we assume rapid buffering such that *w_x_* + *x*_2_ is the slow variable, *w_x_* = *B_x_x*_2_ and 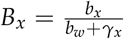, and use the methodology from SI1 and SI2.1 then we have the reduced model

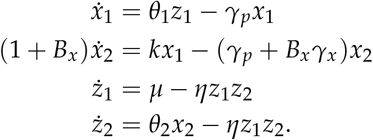

##### SI2.4.a. Steady State Analysis

We next analyse the steady state. We have

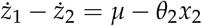

and so the steady state of the output is

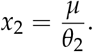

We also have corresponding steady states

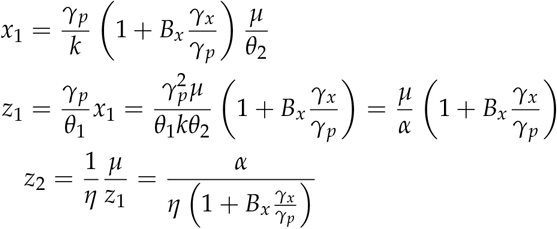

where 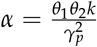.

##### SI2.4.b. Linearised Model

We next analyse the stability of the system. If we linearise about the steady states, we have

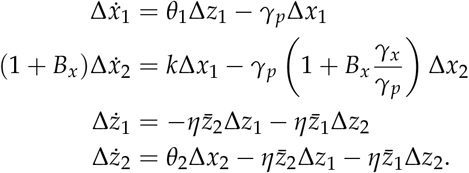

This system can be rewritten as

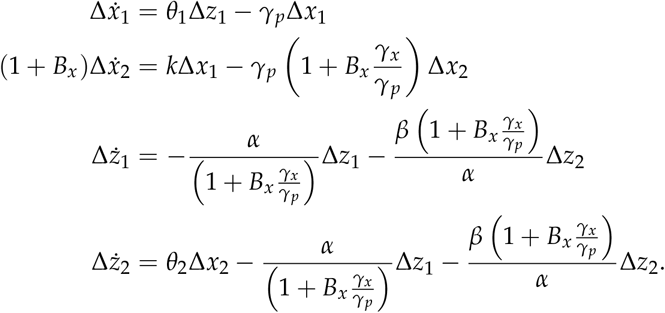

where 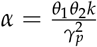 and *β* = *ημ*. Taking the Laplace transforms, we have

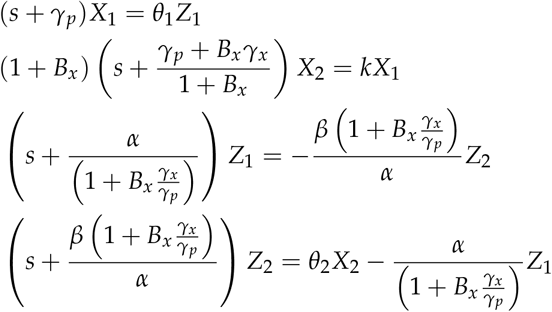

where 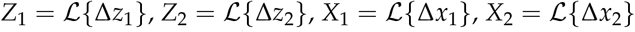 are the Laplace transforms of the time-domain concentration deviations. Substituting, we have

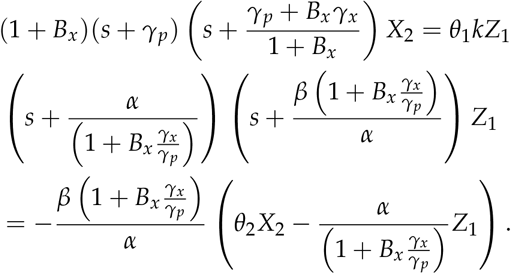

Rewriting and substituting, we have

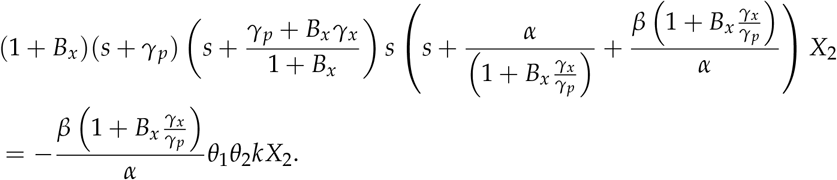

Simplifying and taking the limit of strong binding for the sequestration process in the antithetic integral feedback 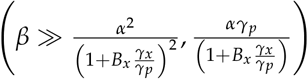 then

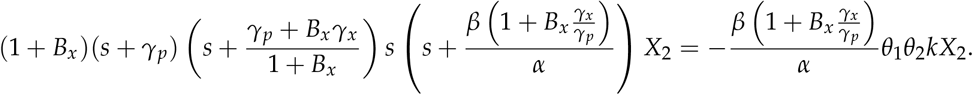

Thus we have the characteristic equation

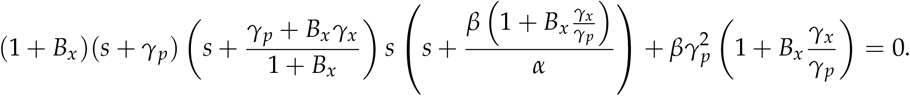

Substituting *s* = *γ_p_σ*, we have

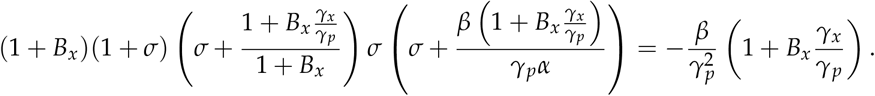

Using the same argument as that in SI2.1, for the stability boundary with strong binding there is a negative real and complex pair of roots in the region 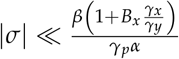, as well as one large negative root.

To determine the boundary of stability, we next determine the conditions for which the roots are purely imaginary. Substituting *s* = *iωγ_p_*, we have

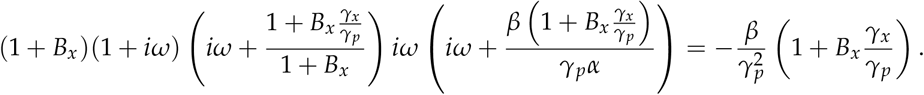

Taking the strong antithetic binding limit where 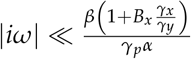, we have

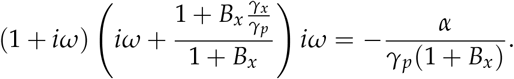

The phase and magnitude constraints are

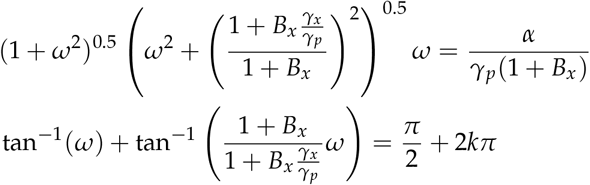

for some integer *k*. Solving the phase constraint, we have

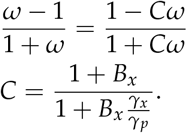

For this constraint we require

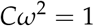

which reduces to

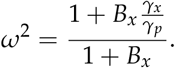

Substituting into the magnitude equation, we have

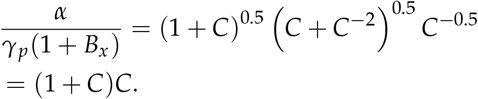

Rearranging, we have the stability condition

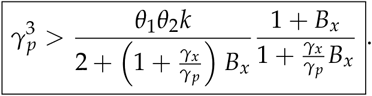

We can differentiate the right hand side with respect to *B_x_* to determine whether increasing *B_x_* has a stabilising effect, If we set

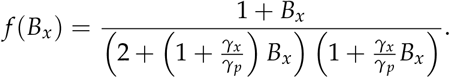

then

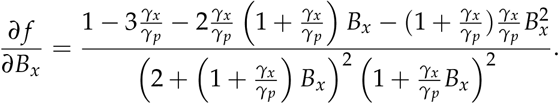

Thus for small *B_x_* then buffering stabilises antithetic integral feedback if 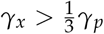, For large *B_x_* then increasing *B_x_* improves stability if *γ_x_* > 0.

#### SI2.5. Stability of Antithetic Feedback: Rapid *x*_1_ Buffering with Degradation

In this section, we analyse the stabilising effect of buffering at the intermediate species *x*_1_ on antithetic integral feedback, where the buffering can be degraded. This section uses identical methodology and obtains equivalent results to SI2.4.

Consider the model (S4) with buffering at *x*_1_ that is degraded

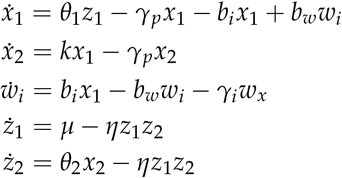

where *w_i_* is the buffering species of *x*_1_, and *b_i_*, *b_w_* are the kinetic rates for the buffering reactions. If we assume rapid buffering such that *w_i_* + *x*_1_ is the slow variable, *w_i_* = *B_i_x*_1_ and 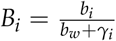, and use the methodology from SI2.4 then

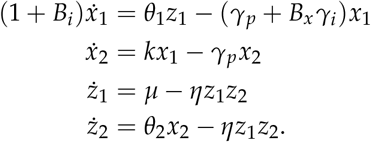

##### SI2.5.a. Steady State Analysis

We next analyse the steady state. We have

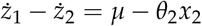

and so the steady state of the output is

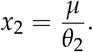

The corresponding steady states of the other species are

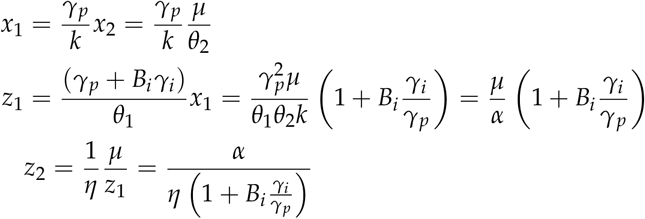

where 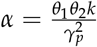.

##### SI2.5.b. Stability Analysis

We next study the stability of the system. If we linearise about the steady states, we have

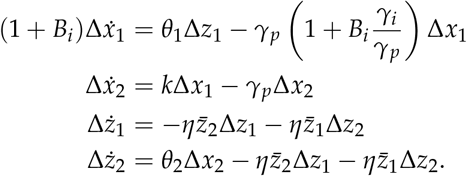

This system can be rewritten as

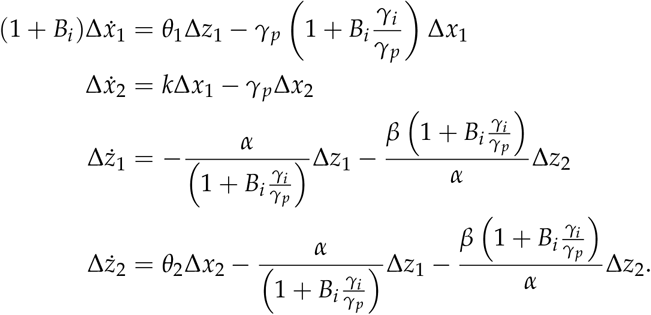

where 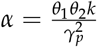 and *β* = *ημ*. Taking the Laplace transforms, we have

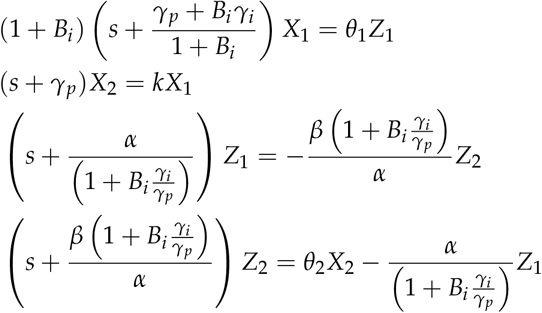

where 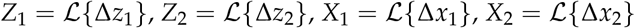 are the Laplace transforms of the time-domain concentration deviations. Substituting, we have

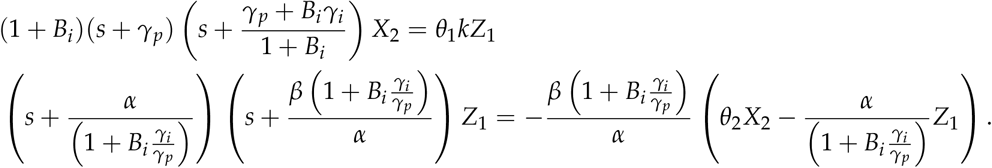

Rewriting and substituting, we have

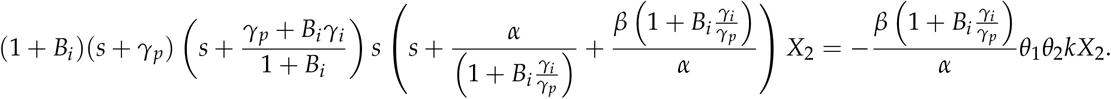

The above equation is equivalent to *x*_2_ buffering (see SI2.5), and so for strong integral binding we have the stability condition

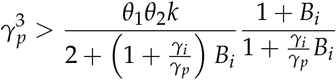

which is equivalent to the case of *x*_2_ buffering.

### SI3. BUFFERING CAN ENABLE NEAR-PERFECT ADAPTATION DESPITE LEAKY INTEGRATION

#### SI3.1. Rapid *x*_2_ Buffering with Degradation can Enable Near-Perfect Adaptation

In this section, we analyse the ability of buffering at *x*_2_ to enable near perfect adaptation by stabilising antithetic integral feedback. Consider the model (S5) with buffering at *x*_2_ that is degraded

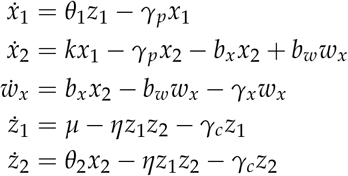

where *w_x_* is the buffering species of *x*_2_, and *b_x_*, *b_w_* are the kinetic rates for the buffering reactions. Assuming that the buffer rapidly reaches equilibrium then using the methodology from SI1 and SI2.1, we have the reduced model

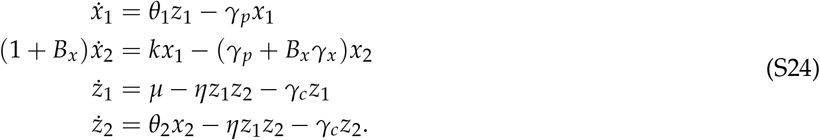

where

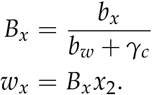

##### SI3.1.a. Steady State Analysis

We next analyse the steady state. We have

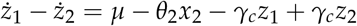

and so the steady state of the output is

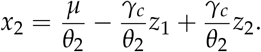

We also have the steady state

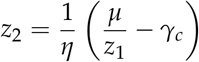

and so

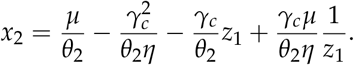

The other species have the steady states

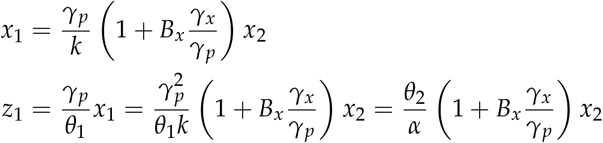

where 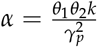. Substituting, we have

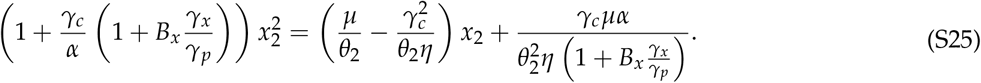

We assume strong binding of the sequestration mechanism in antithetic integral feedback (*η* large), which for steady state is the condition

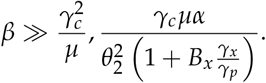

With this assumption, we have

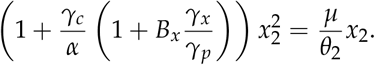

We ignore the zero solution, which corresponds to a negative solution in (S25), and so the steady state concentrations are

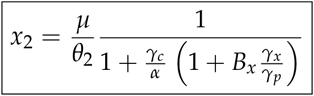

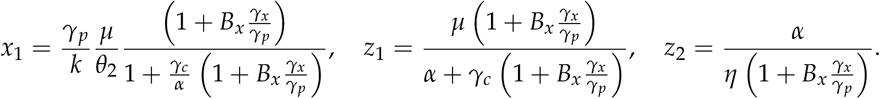

The steady state error of *x*_2_ is

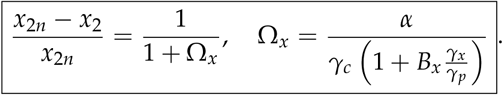

##### SI3.1.b. Stability Analysis

We next study the stability of the system. If we linearise (S24) about the steady state, we have

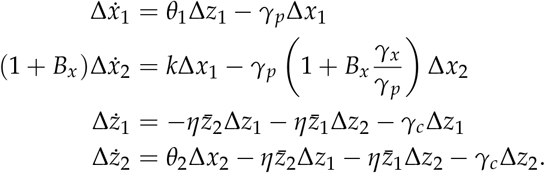

This system can be rewritten as

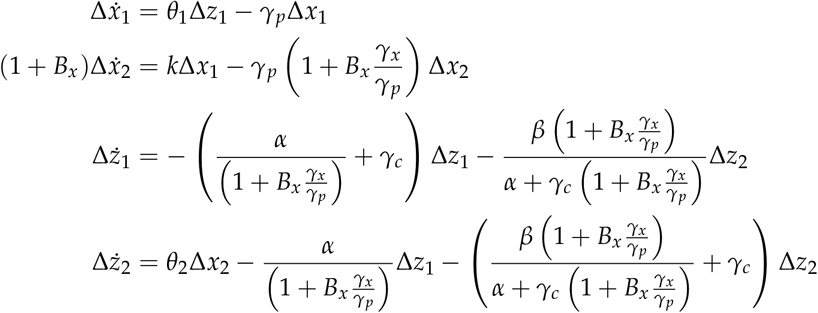

where 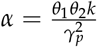 and *β* = *ημ*. Taking the Laplace transforms, we have

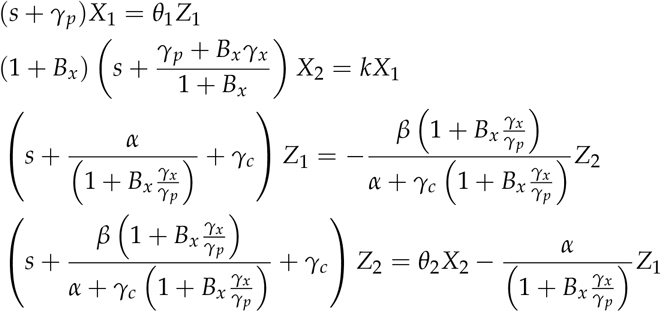

where *X*_1_, *X*_1_, *Z*_1_ and *Z*_2_ are the Laplace transforms of Δ*x*_1_, Δ*x*_2_, Δ*z*_1_ and Δ*z*_2_. Substituting, we have

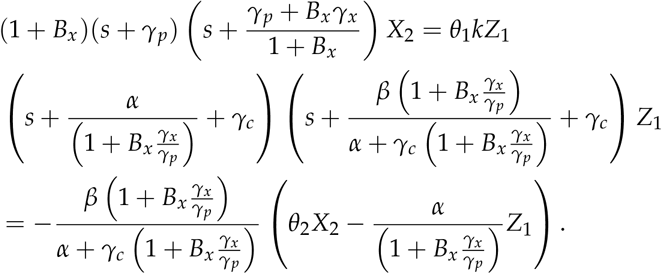

Rewriting and substituting, we have

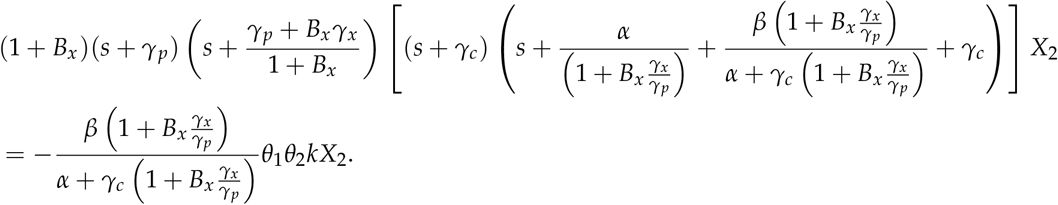

Simplifying and taking the limit of strong binding of the sequestration mechanism in antithetic integral feedback

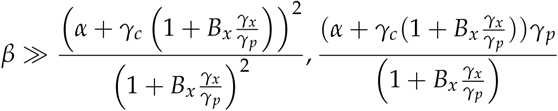

then

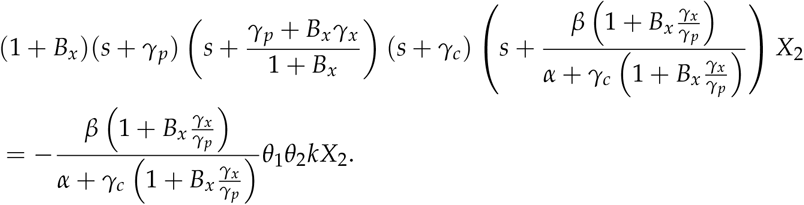

Thus we have the characteristic equation

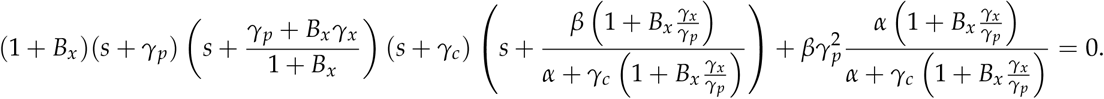

Substituting *s* = *γ_p_σ*, we have

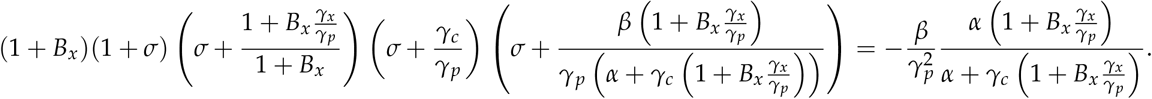

Using the same argument as that used in SI2.1, for the stability boundary with strong binding there is a negative real and complex pair of roots in the region 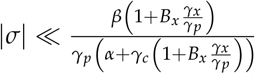, as well as one large negative root.

To determine the boundary of stability, we next determine the conditions for which the roots are purely imaginary. Substituting *s* = *iωγ_p_*, we have

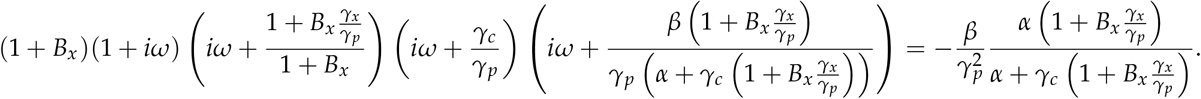

Taking the strong antithetic binding limit where 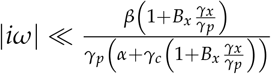, we have

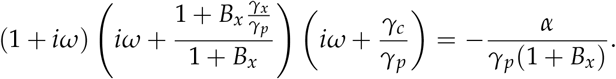

The phase and magnitude constraints are

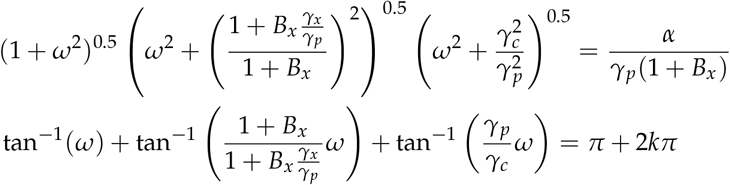

for some integer *k*. Solving the phase constraint, we have

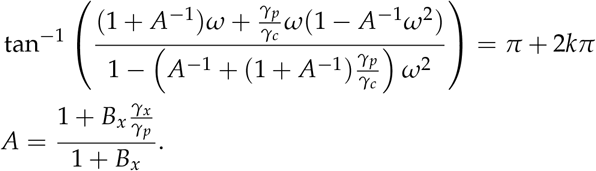

For this we require

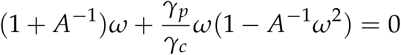

which reduces to

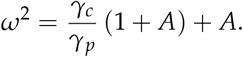

Substituting into the magnitude equation, we have

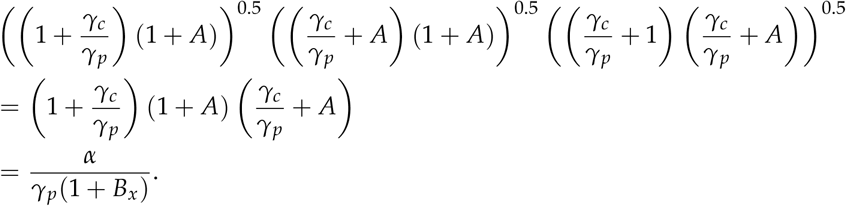

Simplifying, we have

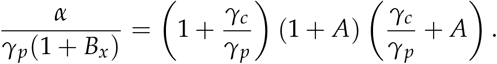

Multiplying by 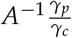, we have

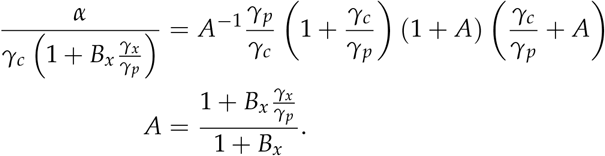

and so the stability condition is

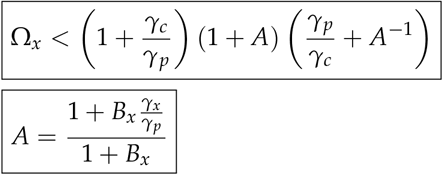

where

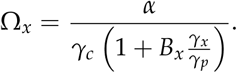

From earlier, we know that the steady state error of *x*_2_ is

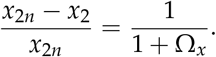

#### SI3.2. Non-Rapid *x*_2_ Buffering can enable Near Perfect Adaptation

In this section, we analyse the ability of non-rapid buffering at *z*_1_ to enable near perfect adaptation by stabilising antithetic integral feedback. We use the model

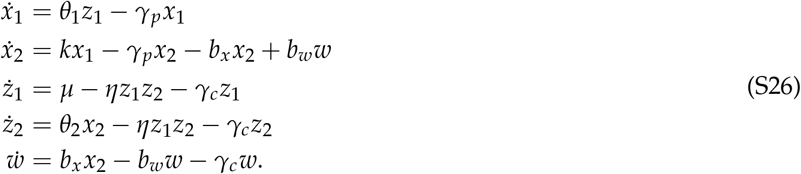

where the buffer *w* is not assumed to rapidly reach equilbrium. The steady state for (S26) is identical to the rapid case in SI3.1 if we replace *γ_x_* by *γ_c_*. The linearisation is

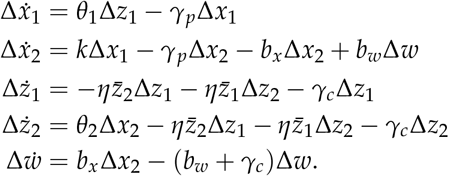

From SI3.1, the steady states are

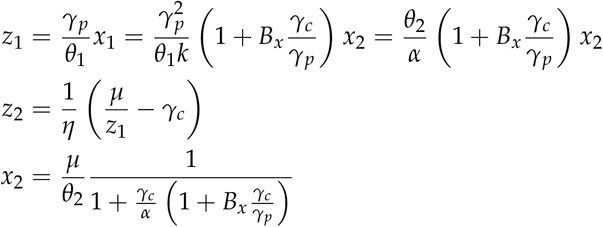

where

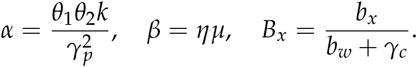

This system can be rewritten

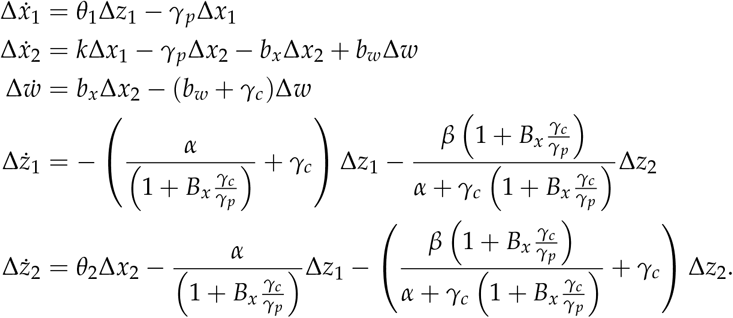

Taking the Laplace transform of 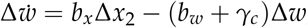, we have

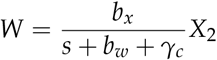

where *W* and *Z*_1_ are the Laplace transforms of *w* and *x*_2_. We have

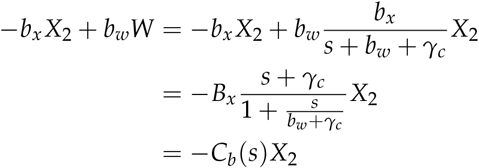

where

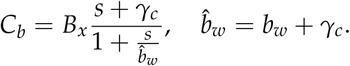

Taking the Laplace transforms, we have

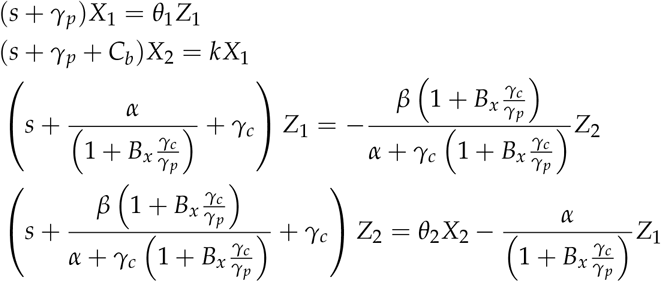

where *X*_1_, *X*_1_, *Z*_1_ and *Z*_2_ are the Laplace transforms of Δ*x*_1_, Δ*x*_2_, Δ*z*_1_ and Δ*z*_2_. Substituting, we have

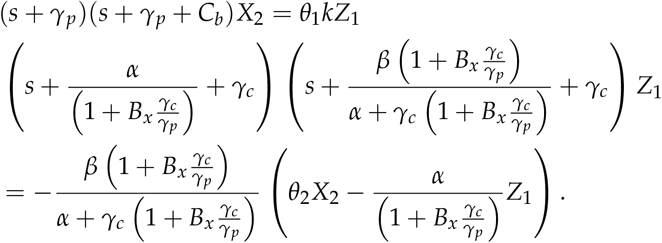

Rewriting and substituting, we have

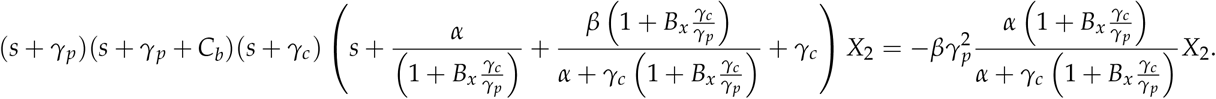

Simplifying and taking the limit of strong binding of the sequestration mechanism in antithetic integral feedback

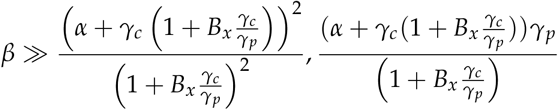

then

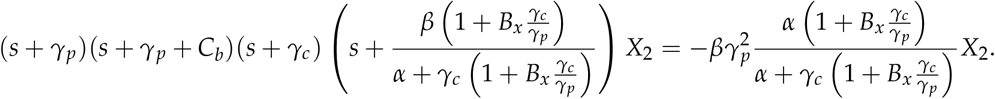

Rewriting *C_b_*, we have

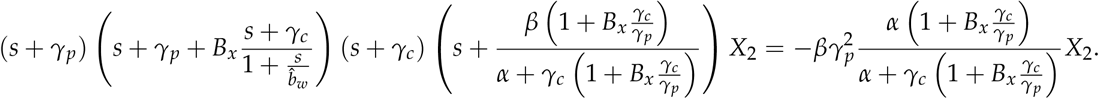

and so we have the characteristic equation

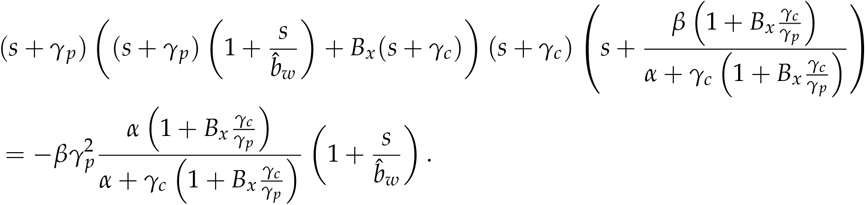

Substituting *s* = *iωγ_p_*, we have

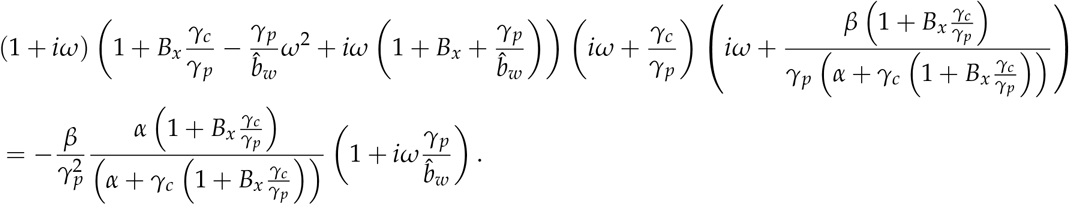

Taking the strong antithetic binding limit where 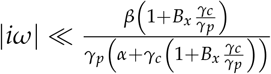, we have

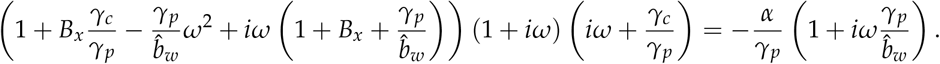

The magnitude constraint is

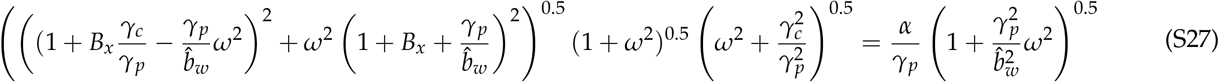

and the phase constraint is

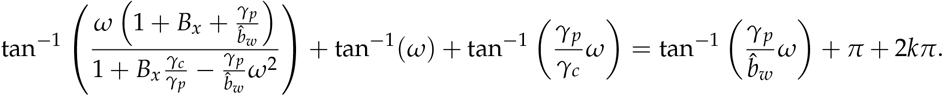

for some integer *k*. Rearranging, we have

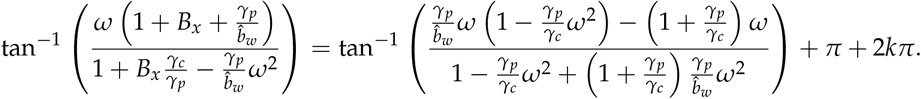

Taking the tangent of both sides and ignoring the trivial solution, we have

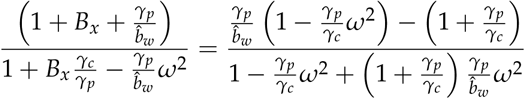

Multiplying out fractions, we have

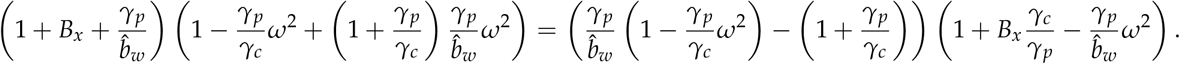

Expanding, we have

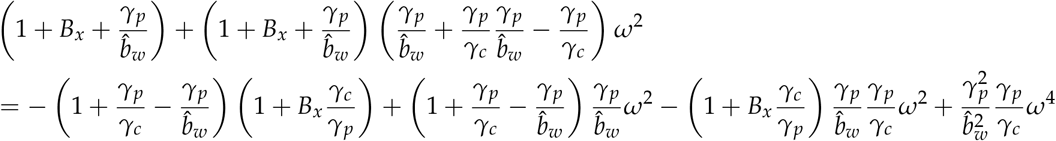

Simplifying, we have

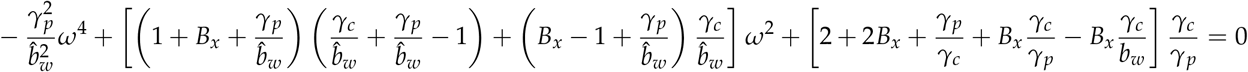

Solving, we have

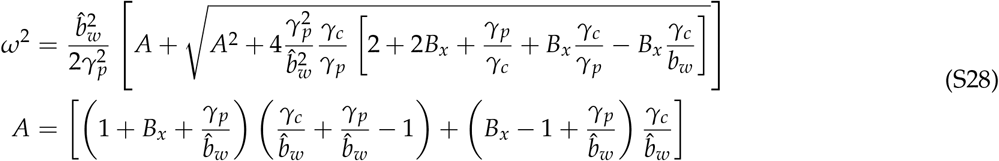

Using the same approach as in previous sections, the magnitude constraint leads to the stability constraint

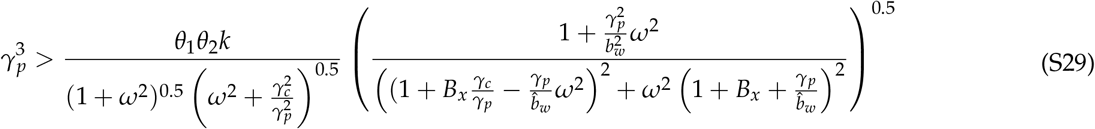

using *ω* from (S28). This can be rewritten

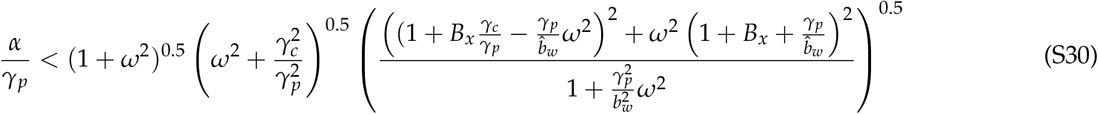

From SI3.1, we have there steady state error

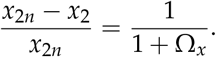

where

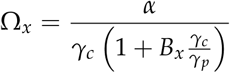

and so the steady state error constraint is

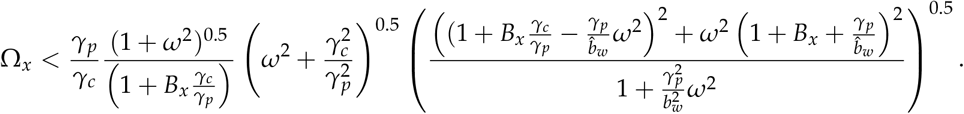

where *ω* is given in (S28).

If we set 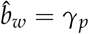 for simplicity then 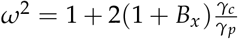 and

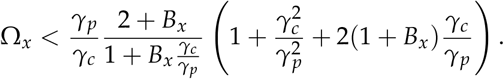

This equation can be rewritten

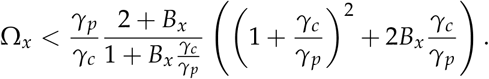

#### SI3.3 Rapid *x*_1_ Buffering with Degradation can enable near-perfect adaptation

In this section, we analyse the ability of buffering at *x*_1_ to enable near perfect adaptation by stabilising antithetic integral feedback. This section uses identical methodology and obtains equivalent results to SI3.1.

Consider the model with *x*_1_ buffering and dilution

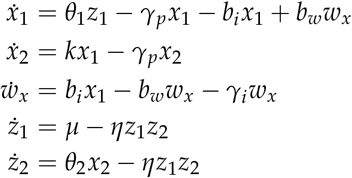

where *w_x_* is the buffering species of *x*_1_, and *b_i_, b_w_* are the kinetic rates for the buffering reactions. Assuming rapid buffering and using the same methodology as previous sections, the reduced model is

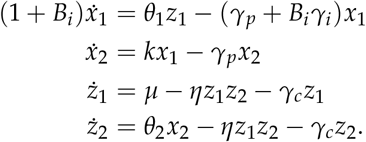

where 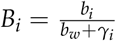 is the buffer equilibrium ratio.

##### SI3.3.a. Steady State Analysis

We next analyse the steady state of the system. We have

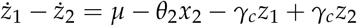

and so the steady state of the output is

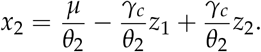

We also have the steady state

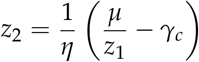

and so

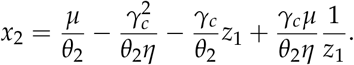

Now at steady state we have

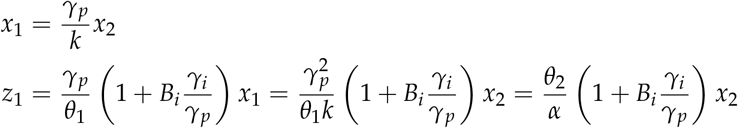

where 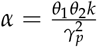. Substituting, we have

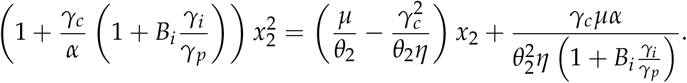

Assuming strong binding

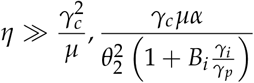

we have

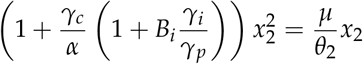

and so, ignoring the solution *x*_1_ = 0, the steady state is

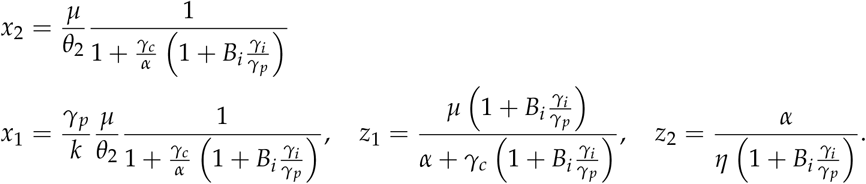

The steady state error of *x*_2_ is

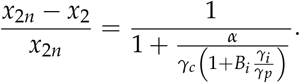

##### SI3.3.b. Stability Analysis

We next study the stability of the system. If we linearise about the steady state, we have

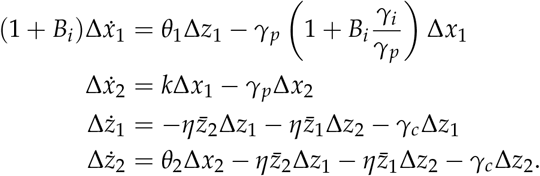

This system can be rewritten as

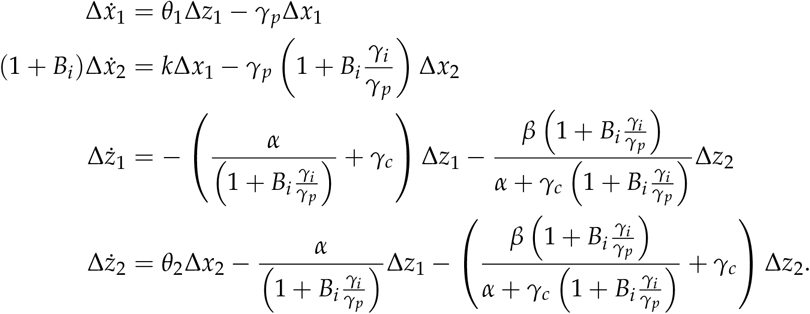

where 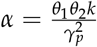 and *β* = *ημ*. Taking the Laplace transforms, we have

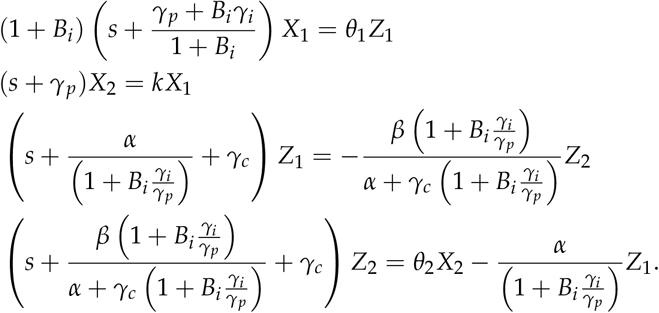

where *X*_1_, *X*_2_, *Z*_1_ and *Z*_2_ are the Laplace transforms for Δ*x*_1_, Δ*x*_2_, Δ*z*_1_ and Δ*z*_2_. Substituting, we have

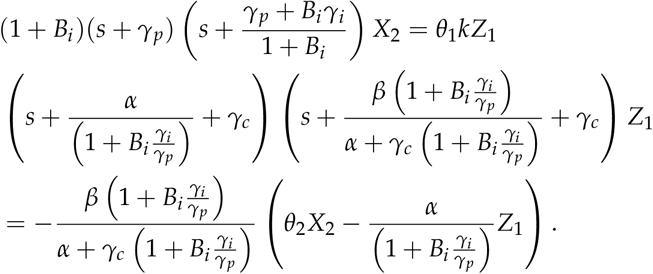

Rewriting and substituting, we have

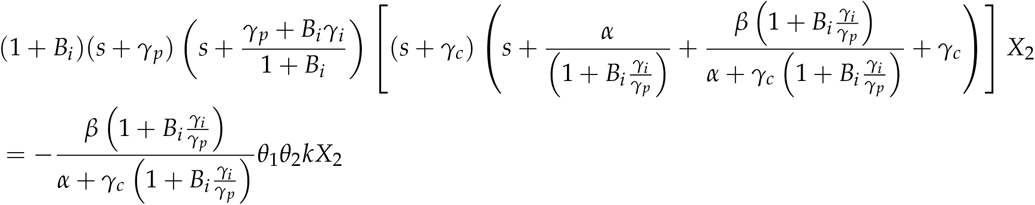

The above equation is equivalent to that for *x*_2_ buffering (see SI3.1), and so for strong integral binding we have the equivalent stability constraint and steady state error

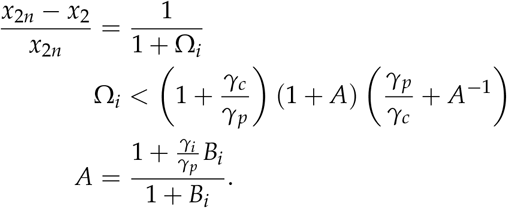

#### SI3.4. Rapid *z*_1_ Buffering with Degradation has a trade-off due to leaky integration

In this section, we analyse the trade-offs for rapid buffering at *z*_1_ on stability and the steady state error from perfect adaptation. For buffering at *z*_1_ with dilution, we use the model

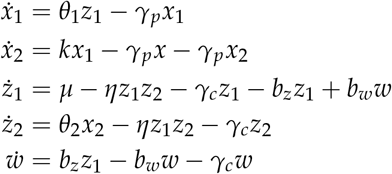

where *x*_2_ is the output concentration being controlled, *x*_1_ is another concentration in the process being controlled, and *z*_1_ and *z*_2_ represent the molecular species involved in the perfect adaptation mechanism. Assuming that the buffer is rapid then *w* is at quasi-steady state then

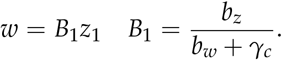

If *x_T_* = *w* + *z*_1_ is the slow variable then *x_T_* = (1 + *B*_1_)*z*_1_. Thus *ẋ_T_* = (1 + *B*_1_)*ż*_1_ and so

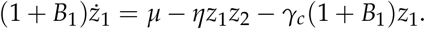

Thus we have

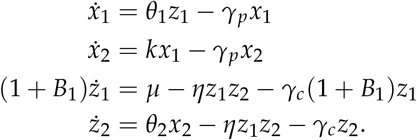

##### SI3.4.a. Steady State Analysis

We next determine the steady state and and any error from perfect adaptation. For the case of dilution, we have

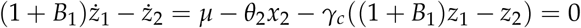

resulting in

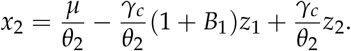

We also have

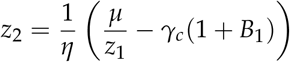

and so

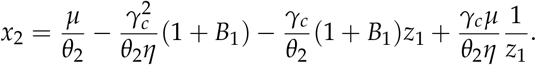

Now at steady state we have

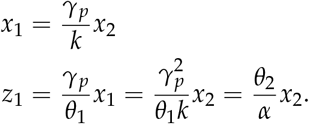

where 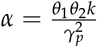. Substituting, we have

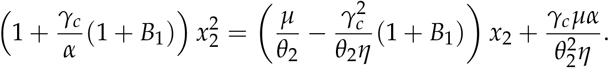

Assuming strong binding of the sequestration mechanism

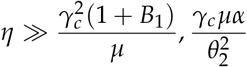

we have

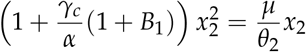

and so, ignoring the zero solution, the steady state is

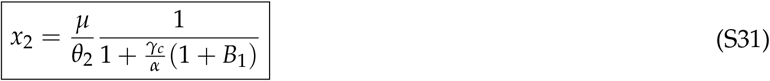

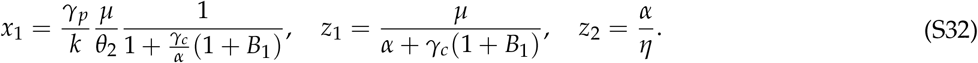

The steady state error of *x*_2_ is

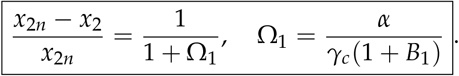

We can see that increasing *B*_1_ increases the steady state error of *x*_2_ when there is degradation/dilution of *z*_1_ and *z*_2_.

##### SI3.4.b. Stability Analysis

We next study the stability of the system with degradation. If we linearise about the steady states, we have

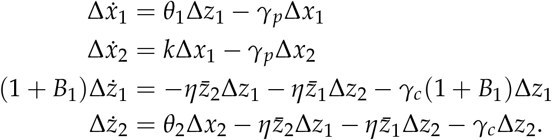

This system can be rewritten as

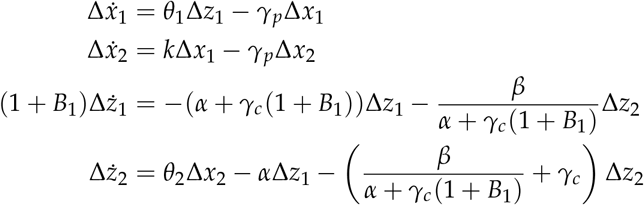

where

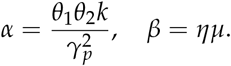

Taking the Laplace transforms, we have

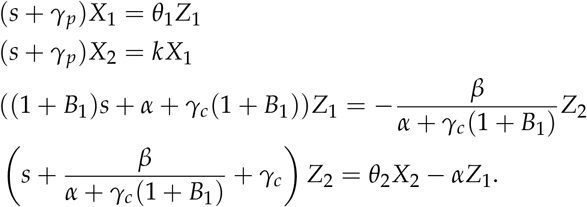

where *X*_1_, *X*_2_, *Z*_1_ and *Z*_2_ are the laplace transforms for Δ*x*_1_, Δ*x*_2_, Δ*z*_1_ and Δ*z*_2_. Substituting, we have

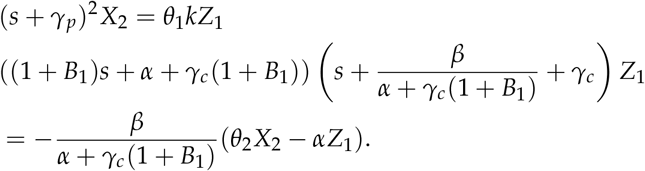

Rewriting and substituting, we have

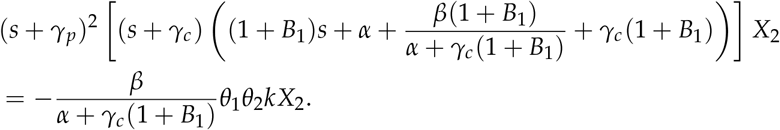

Taking the limit of strong binding 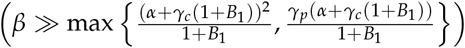 then

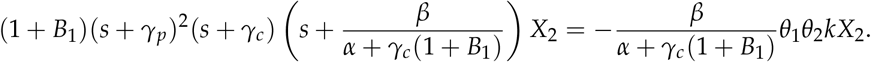

Thus we have the characteristic equation

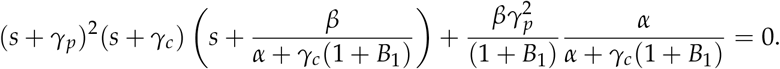

Substituting *s* = *γ_p_σ*, we have

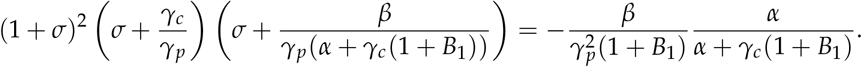

Using the same argument as above, for the stability boundary with strong binding there is a negative real and complex pair of roots in the region 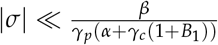, as well as one large negative root.

To determine the boundary of stability, we next determine the conditions for which the roots are purely imaginary. Substituting *s* = *iωγ_p_*, we have

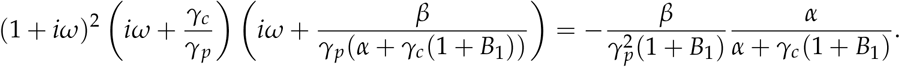

Taking the strong antithetic binding limit where 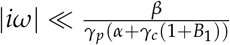, we have

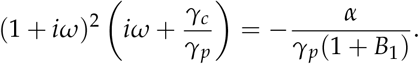

The phase and magnitude constraints are

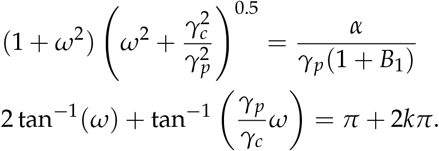

for some integer *k*. Solving the phase constraint, we have

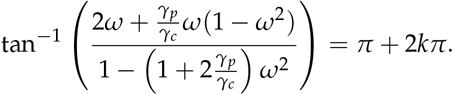

For this, we require

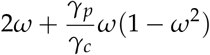

which reduces to

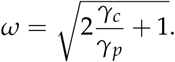

Substituting into the magnitude equation, we have

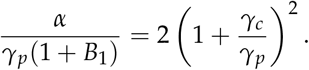

As a consequence, the stability constraint is

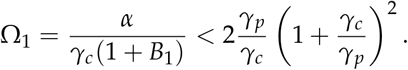

We can observe that increasing *B*_1_ improves the stability constraint. However, the steady state error of *x*_2_ is

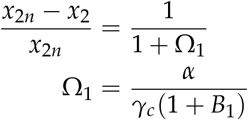

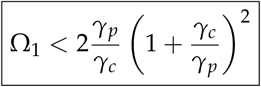

Thus there is a steady state error constraint that is independent of *B*_1_, and so increasing *B*_1_ does not enable the removal of leaky integration.

#### SI3.5. Non-Rapid *z*_1_ Buffering can allow Near Perfect Adaptation

In this section, we analyse the ability of non-rapid buffering at *z*_1_ to enable near perfect adaptation by stabilising antithetic integral feedback. We use the model

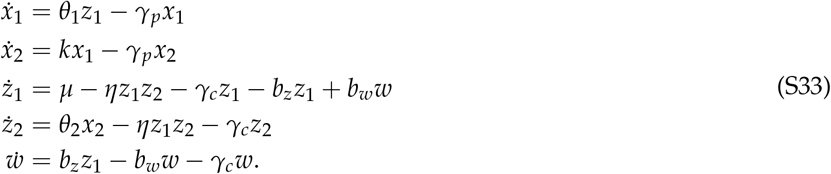

where the buffer *w* is not assumed to rapidly reach equilibrium. The steady state for (S33) is identical to the rapid case in SI3.4. The linearisation is

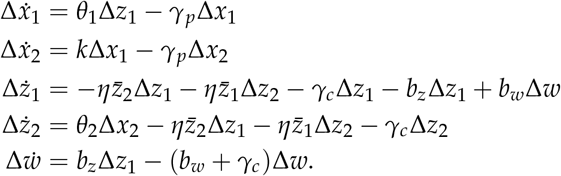

This system can be rewritten

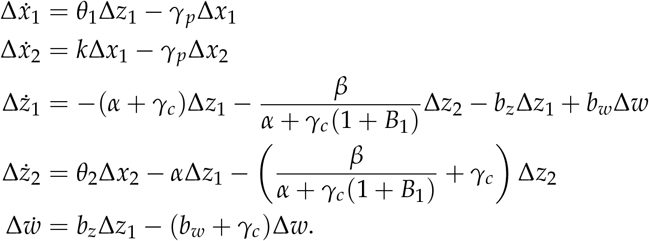

where

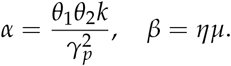

Taking the Laplace transform of 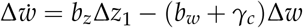, we have

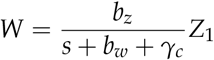

where *W* and *Z*_1_ are the Laplace transforms of *w* and *z*_1_. We have

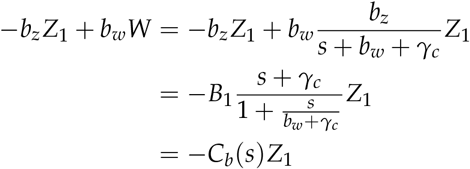

where

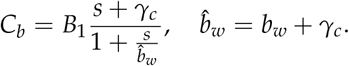

Thus

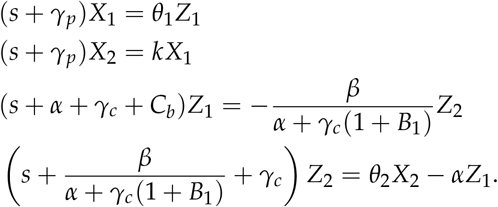

Combining, we have

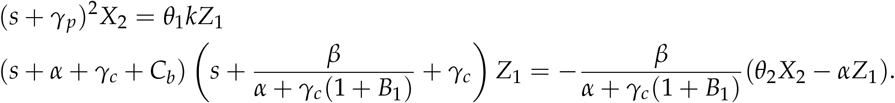

Simplifying, we have

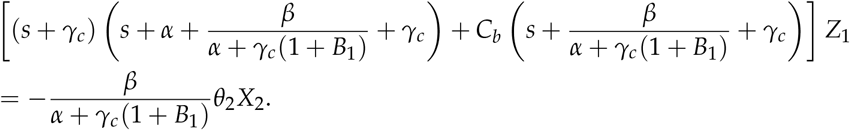

Taking the strong antithetic binding limit of the sequestration mechanism (*β* ≫ max{(*α* + *γ_c_*)(*α* + *γ_c_*(1 + *B*_1_)), *γ_p_*(*α* + *γ_c_*(1 + *B*_1_))}), we have

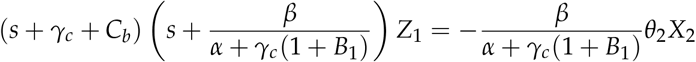

and so

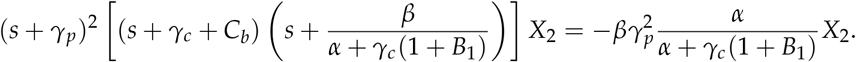

Rewriting *C_b_*, we have

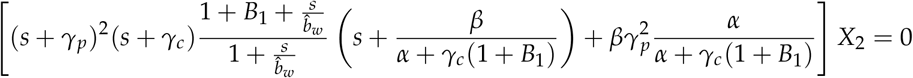

or

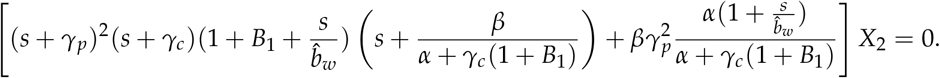

Substituting *s* = *iωγ_p_*, we have

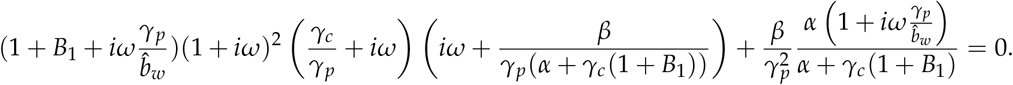

Taking the strong antithetic binding limit where 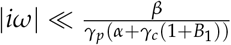, we have

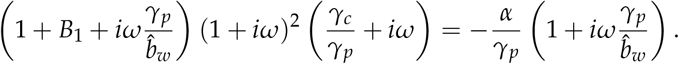

The magnitude constraint is

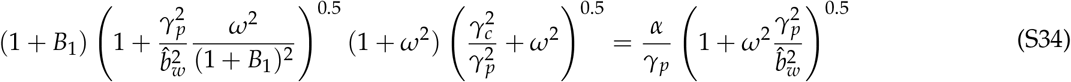

and the phase constraint is

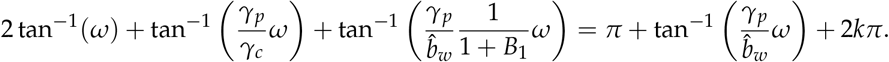

for some integer *k*. Using trigonometric identities, we have

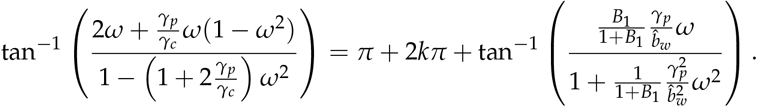

This can be simplified to

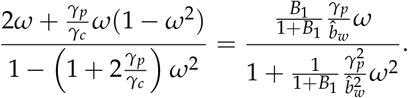

Ignoring the trivial solution *ω* = 0, we have

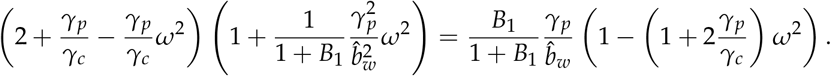

Rewriting, we have

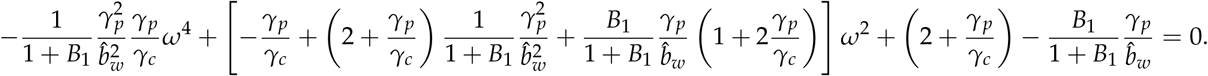

Simplifying, we have

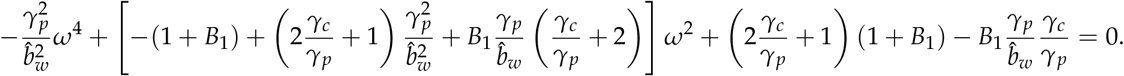

Solving, we have

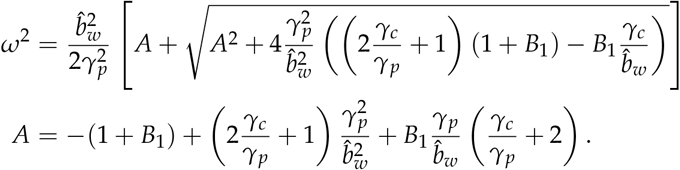

From the magnitude constraint S34, we have the stability constraint

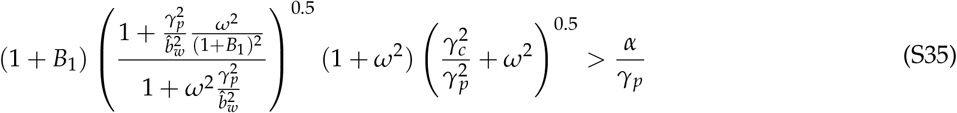

which can be rewritten in terms of steady state error 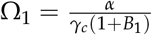 as

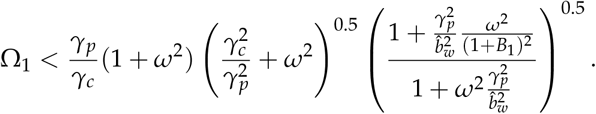

For 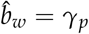, we have

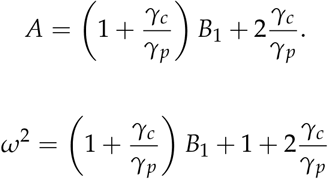

and the stability constraint

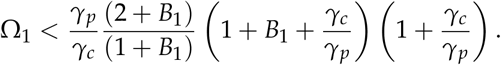

### SI4. CHARACTERISATION OF ANTITHETIC FEEDBACK WITH BUFFERING

In this section we characterise the feedback in the system with antithetic feedback and buffering of the control species *z*_1_, *z*_2_.

#### SI4.1. Characterisation with Rapid Buffering

We first characterise antithetic feedback with rapid buffering. We use the model

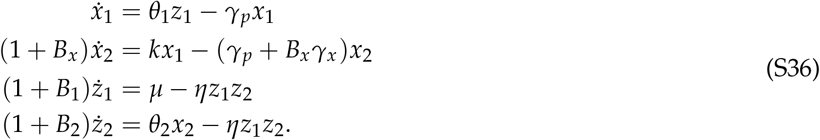

It can be observed that rapid buffers of the controller change the time-scale for *z*_1_ and *z*_2_. In integral feedback, the time-scale is inversely proportional to gain, and this relationship can be observed in the following characterisation of integral gain. We have

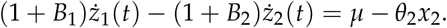

Integrating, we have

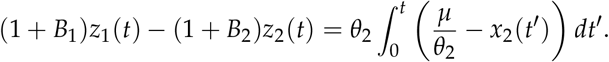

Rearranging, we can observe that the feedback control input *u* into the plant (i.e. (*x*_1_, *x*_2_) subsystem) is

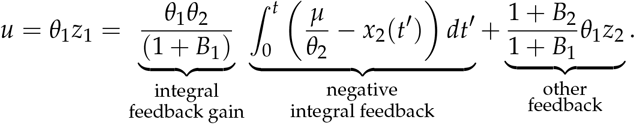

We can observe an integral feedback term and a second feedback term that is dependent upon *z*_2_. The other feedback term becomes negligible and the integral term dominates the overall feedback if

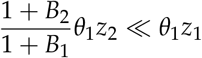

which can occurs if 1 + *B*_1_ is sufficiently large or if *z*_2_ is sufficiently small. The latter can be achieved via large *η*, which drives *z*_2_ to a much lower concentration than *z*_1_. For this case

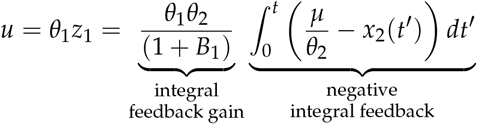

where we can observe that the feedback gain is dependent on *B*_1_.

#### SI4.2. Characterisation with Non-rapid *z*_1_ Buffering

For the case of non-rapid *z*_1_ buffering, we have

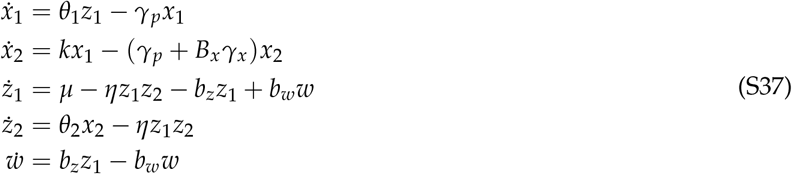

We have

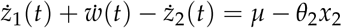

where the integral is

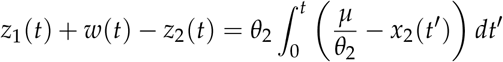

and so perfect adaptation occurs. Rearranging, we have the feedback

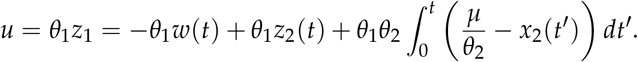

Taking Laplace transforms, we have

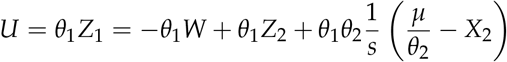

where *Z*_1_, *W*, *Z*_2_, *X*_2_ are the Laplace transforms of *z*_1_, *w*, *z*_2_, *x*_2_. Taking the Laplace transforms of (S37), we also have

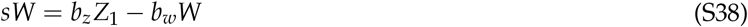

and so

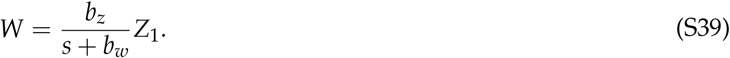

Substituting, we have

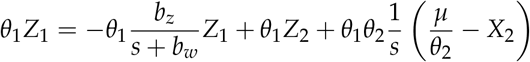

and so the control input is

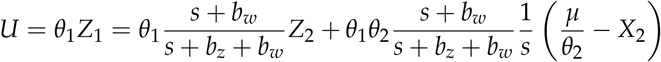

If *Z*_2_ is small due to large *η*, then we have

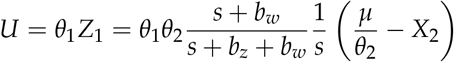

Thus for non-rapid buffering (small *b_w_*) we are able to add a zero and pole and zero to the controller, where the zero is small than the pole. This result is equivalent to the integral controller in series with a lead controller, where the latter is know to help stabilise systems^1^.

We can see the effect of non-rapid buffering on the integral component of feedback gain separately by determining the gain of the control input in the asymptote as *s* → 0, where we have

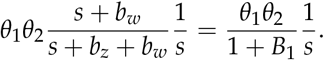

Thus the integral component of feedback gain is 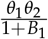, which is identical to the rapid case.

#### SI4.3. Bode Integral of Buffering and Antithetic Integral Feedback

Feedback is a highly effective method of robust regulation, but this mechanism is subject to fundamental limits. The Bode integral describes one of these fundamental limit, where improving the regulation at one frequency of a disturbance will worsen regulation of disturbances at other frequencies. Here, we show that topology 3 has the ability to reduce the fundamental limit on feedback and thus uniformly improve output regulation at all frequencies, while topology 1 does not. While we show the two state example from above to illustrate the concept, the Fourier transform description below is written generally as the Bode Integral holds for more general processes controlled by antithetic feedback and buffering.

We use the example model of the system

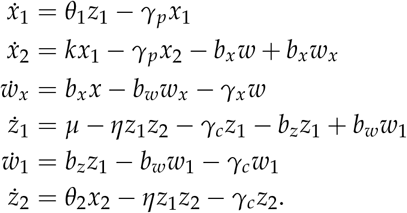

We can write the linearised model in an open-loop form with the two states

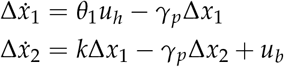

where 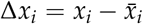 for *i* = 1,2, 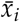 is the steady state of *x_i_* for the closed loop system, *u_h_* is the process input for antithetic integral feedback and *u_b_* is the input for buffering at *x*_2_. Buffering at *z*_1_ and antithetic integral feedback act through the input *u_a_* while buffering at *x*_2_ acts through *u_b_*.

If we take the Fourier transform, we can describe the open loop model in general terms by^3^

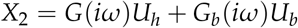

where *X*_2_, *U_h_, U_b_* are the Fourier transforms of *x*_2_, *u_h_*, *u_b_* respectively, *G* is the transfer function from *U_h_* to the output *X*_1_ and *G_b_* is the transfer function from *U_h_* to *X*_1_. With two control inputs we have the loop transfer function^3^

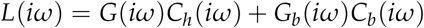

where *C_h_* is the transfer function for the antithetic feedback controller (i.e. from *X*_2_ to *U_h_*) and *C_b_* is the transfer function for the output buffer (i.e. from *X*_2_ to *U_b_*). The sensitivity function quantifies the improvement or worsening in regulation at each frequency for the ‘closed-loop’ system, where a smaller magnitude implies improved regulation, and can be described by

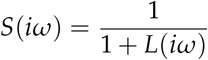

In this case the sensitivity function incorporates the regulatory effect of both feedback and buffering^3^.

The Bode integral is a fundamental constraint on the effectiveness of feedback in any system. It provides a constraint on the overall regulatory effectiveness in terms of the sensitivity function. If the system without feedback is stable then Bode’s integral with output buffering (Topology 3) is^3^

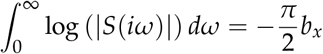

where the integral of *S*(*iω*) represents an overall measure of regulation and *b_x_* is the kinetic rate of the forward buffering reaction. The integral of *S*(*iω*) sums the effect of oscillating disturbances at different frequencies *ω*. Without buffering, if regulation is improved at one frequency of regulation, it worsens at other frequencies. However, increasing *b_x_* reduces the whole integral. Thus buffering can uniformly improve regulation and thus improve the trade-off.

In contrast, control species buffering (Topology 1) is part of the feedback regulation mechanism using the same control input and so Bode’s integral is^3^

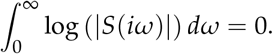

Thus control buffering does not remove fundamental constraints, despite stabilising buffering. The tradeoff remains such that improving regulation at one frequency will worsen regulation of disturbances at other frequencies.

